# Irregular nucleosome positioning governs a crystalline to liquid-like phase transition and tunes chromatin accessibility

**DOI:** 10.64898/2026.08.24.746683

**Authors:** Waad AlBawardi, Matthew Thomas, Gert-Jan Kuijntjes, Hannah Wheeldon, Artur Kaczmarczyk, James Allan, James Ding, Michael Chiang, Willem Vanderlinden, John van Noort, Davide Marenduzzo, Chris Brackley, Nick Gilbert

## Abstract

Chromatin must fold tightly enough to protect the genome while being sufficiently accessible for DNA dependent processes such as transcription. The physical rules that balance these competing roles remain unclear, as DNA sequence encodes both biochemical information such as transcription factor binding sites, and biophysical cues that shape chromatin structure. Here, using synthetic chromatin fibres assembled from physiologically relevant DNA sequences, we show that nucleosome positioning dictates the material state of chromatin. Heterochromatin-like sequences produce compact fibres stabilised by nucleosome stacking, whereas euchromatin-like sequences generate irregular nucleosome positioning that yields disrupted, heterogeneous, and mechanically deformable fibres. Quantitative polymer modelling reveals that these irregular arrays are highly dynamic, continually sampling a broad ensemble of conformations as nucleosome stacking breaks down. We identify two previously unrecognised thresholds encoded by nucleosome positioning: minimal positional irregularity (2–3 bp) triggers a transition from an ordered paracrystalline state to a liquid-like phase, whereas an order of magnitude greater irregularity (∼18 bp) is required to generate accessibility and mechanical fragility permissive for transcription factor binding. Euchromatin-like arrays reside at this accessibility threshold. These findings indicate that nucleosome positioning tunes chromatin toward or away from critical structural states that couple genome protection, chromatin dynamics, and transcriptional potential—providing a physical mechanism that helps connect DNA sequence to gene expression.

## Introduction

In higher eukaryotes chromatin protects the genetic code from environmental and cellular insults yet must allow rapid access to the underlying DNA for essential nuclear processes such as transcription^1–4^. Understanding how regulatory factor binding (a biochemical behaviour) operates in the context of chromatin structure (a biophysical property) is challenging because chromatin forms a complex polymer in the nucleus^5^, and factor binding occurs within this crowded environment^6^. These constraints make direct investigation of chromatin folding difficult, necessitating robust *in vitro* models^7^.

Synthetic chromatin fibres are typically made from purified histone octamers, linker histones and a DNA template with high-affinity nucleosome binding sites such as the artificial ‘Widom 601’ or Xenopus 5S sequences. Chromatin reconstituted onto this substrate using a canonical 197 bp nucleosome repeat forms fibres that have a diameter of 34 nm and are condensed^7^, analogous to satellite-containing heterochromatin^8^. This order depends on nucleosome interactions, which can be described biophysically as paracrystalline, as the nucleosome axis defines a local direction associated with orientational order^9^.

However, microscopic imaging suggests there are few ordered chromatin fibres in nuclei and instead they are disrupted with diameters that vary between 4 and 24 nm^10,11^. Furthermore, small-angle X-ray scattering (SAXS) of chromatin in solution indicates that mitotic chromatin lacks a periodic structure beyond the 11 nm scale^12^, raising the possibility that fibres can interdigitate forming a polymer-melt^13,14^, in a way that is incompatible with the ordered structures found with ‘Widom 601’ arrays.

Although ‘Widom 601’ derived fibres are suitable for structural studies due to their regularity, they are not representative of bulk native euchromatin, which contains DNA sequences of differing affinities, and nucleosomes positioned with variable spacing^15–18^. Single-cell and long-read mapping approaches describe variable positioning patterns in different genomic regions. Constitutive heterochromatin domains show regular spacing between nucleosomes, while being poorly phased. In contrast, active genes have a non-uniform nucleosome distribution along single molecules, but are strongly phased^17,19,20^. Recent work has explored how linker DNA length modulates inter- and intra-fibre associations within regular arrays^21,22^, but how nucleosome array irregularity, which is prevalent in euchromatin, affects chromatin biochemical and biophysical properties remains unclear^10–14^. Standard chromatin reconstitution procedures^23^ are well-suited to high affinity ‘Widom 601’ DNA arrays but are not optimal for DNA sequences with heterogenous nucleosome binding sites, that are typically observed in euchromatin. To investigate physiologically relevant arrays, we developed an alternative approach for reconstituting synthetic chromatin using an excess of nucleosome core particles that resulted in fully saturated chromatin fibres, featuring nucleosome densities and linker DNA lengths that reflect chromatin assembled in cells^24^.

We found that chromatin fibre folding was primarily determined by nucleosome positioning. Regularly spaced nucleosomes formed a paracrystalline structure, which transitioned into a more disordered, liquid-like heterogeneous state—resembling the nematic-to-isotropic phase transition described by the Maier-Saupe theory for liquid crystals^25^. Utilising a coarse-grained model for chromatin, that we term NucHOP (Nucleosome-resolved Higher order Polymer), we quantified chromatin fibre folding. We found that euchromatic fibres are far from a thermodynamically stable state and instead sequence-encoded nucleosome positioning can influence chromatin fibre accessibility, mechanical stability and dynamics.

## Results

### A synthetic *in vitro* model of euchromatin fibres

To understand the physical rules that relate the different biochemical and biophysical layers of information encoded in the DNA sequence we constructed novel DNA templates that we predicted would position nucleosomes irregularly, analogous to positioning patterns observed in euchromatin^17^. We reconstituted two non-repetitive arrays using nucleosome positioning sequences selected from the ovine β-lactoglobulin (*BLG*) gene^26^ (Extended data 1). As the *BLG* locus also encompassed regions that lack nucleosome positioning signals, 197 bp nucleosome binding sequences were daisy chained to exclude large nucleosome free regions and allow comparison of arrays of the same size and nucleosome number. Reconstitution 1 (R1) was composed of 25 unique 197 bp nucleosome binding sites, and Reconstitution 2 (R2) consisted of 12 unique *BLG* nucleosome positioning sequences interspersed with the ‘Widom 601’ positioning motif, to create a partially repetitive array. The strongly positioning Widom-601 template (25 x 197 bp) was used to model heterochromatin-like structures (Fig 1a).

**Figure 1.**
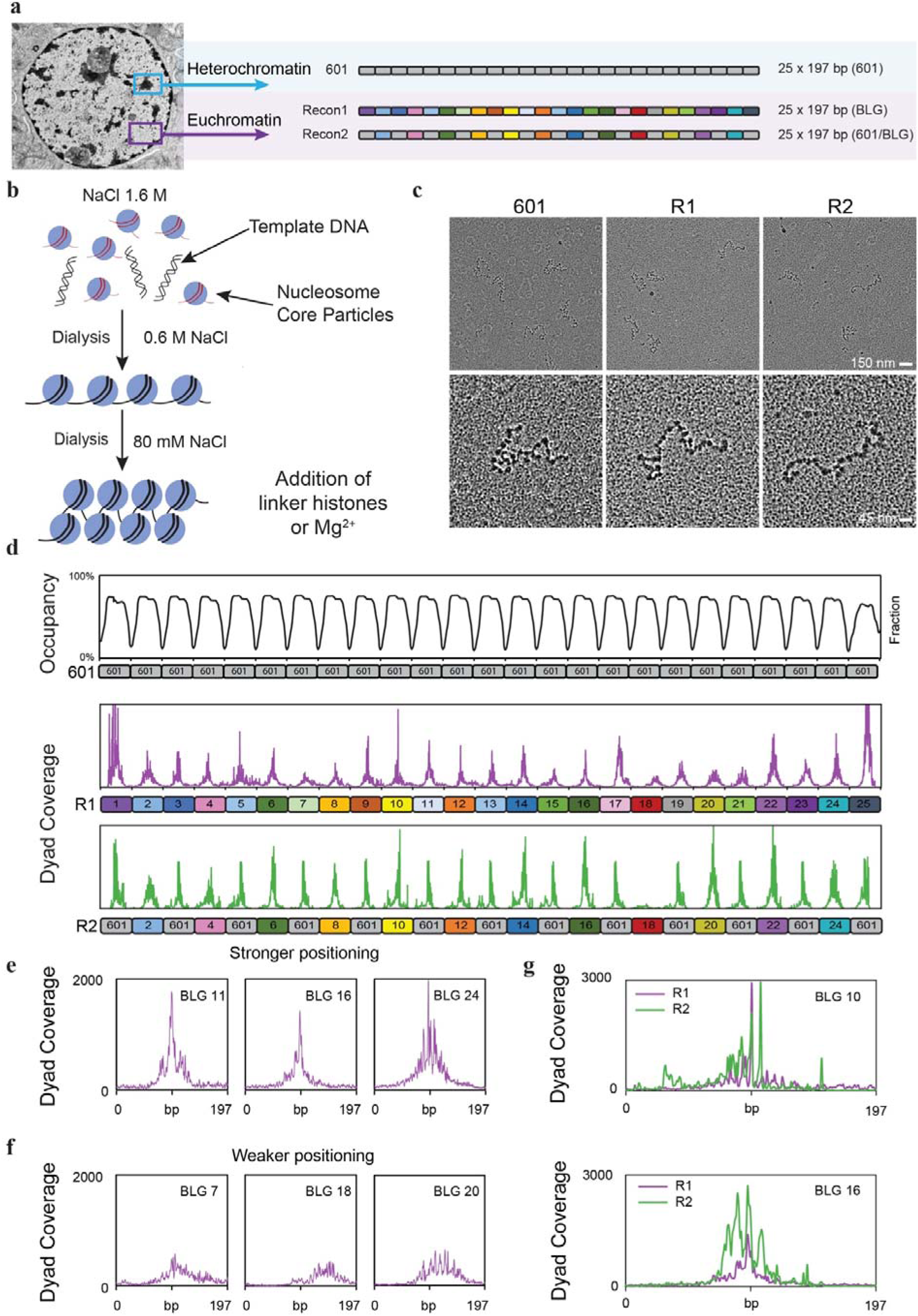
Synthetic chromatin fibre reconstituted using NCPs on physiologically relevant DNA templates. **a,** Schematic of synthetic chromatin fibre templates used in this study. The Widom 601 template is composed of 25 × 197 bp 601 sequences (shown in light grey). Recon 1 contains 25 unique *BLG* positioning sites (depicted in unique colours). In the Recon 2 template each odd-numbered site is replaced by a 601-positioning sequence (shown in light grey). **b**, Schematic showing chromatin reconstitution by histone transfer. Nucleosome core particles (NCPs) are the donor source of histones which are transferred onto template DNA by salt dialysis, and the subsequent addition of linker histones (or Mg^2+^) to facilitate physiological fibre folding. **c,** Representative TEM images showing nucleosome arrays reconstituted via histone-transfer at a 20:1 NCP:template ratio. After reconstitution, fibres were dialysed into low salt (1 mM NaCl) giving unfolded arrays. **d,** Nucleosome position across template DNAs. Top: Nucleosome occupancy over the Widom 601 template derived from Fiber-seq^31^. Bottom: Dyad coverage maps over the R1 and R2 templates derived from MNase-seq data. **e,** Examples of strong nucleosome positioning sites on the R1 template, coincident with BLG 11, 16 and 24. **f,** Examples of weak nucleosome positioning at BLG sites 7, 18 and 20 on the R1 template. **g,** Dyad coverage at BLG 10 and BLG16 are different in R1 and R2 are due to different flanking nucleosomes.

Conventional chromatin reconstitution approaches necessitate precise titration of core histone octamers (CHOs) to achieve correct nucleosome loading^23^. Initial reconstitution attempts using salt dialysis with purified CHOs in the presence of competitor DNA yielded fibres with undersaturated and inconsistent nucleosome occupancy. To correctly load nucleosomes, we modified a histone transfer reconstitution method^24^. Template DNA was combined with excess donor chicken erythrocyte nucleosome core particles (CE NCPs) in high salt concentrations and dialysed down to physiological salt conditions over several hours to facilitate NCP binding at energetically favourable sites (Fig 1b). Reconstitution with a 20:1 NCP to template ratio, by mass, gave complete nucleosome loading (Extended data 2a), which was confirmed by reconstitution of low affinity nucleosome binding sites (Extended data 2b-c), and electron microscopy verified homogeneous reconstitution across templates (Fig 1c).

To characterise nucleosome positioning in the novel templates, we conducted micrococcal nuclease digestion with sequencing (MNase-seq) (Fig 1d) to reveal a striking difference in the nucleosome positioning patterns of 601 and non-repetitive templates. Whilst nucleosomes in 601 arrays occupy precise positions^27^ (Extended data 3a), as observed in heterochromatin^28,29^, the nucleosome spacing in R1 and R2 was more irregular, with measured spacings of 50.1 ± 28.9 bp and 49.6 ± 24 bp, respectively (Fig 1d). Examination of the mean dyad distribution within non-repetitive templates showed that 601 nucleosomes (present in R2 fibres) have distinct dyad peaks, with similar binding affinities, while *BLG* dyads distribute more broadly along the positioning sites, reflecting weaker phasing (Fig 1d). Well-positioned *BLG* nucleosomes include *BLG* 11, 16 and 24, which have sharp dyad coverage peaks (Fig 1e). In contrast, *BLG* 7, 18 and 20 have a broad distribution of mapped dyads suggesting weaker positioning (Fig 1f), representative of the different affinities for nucleosomes across the genome. The position of nucleosomes at *BLG* sites was modulated by flanking ‘601’ sites, clearly observed by a shift in nucleosome position for *BLG* 10 and *BLG* 16 between the R1 and R2 templates (Fig 1g).

To examine the heterogeneity of nucleosome binding in the population of chromatin fibres we used fibre-seq (Extended data 3)^30,31^. 601 fibres showed highly regular and homogenous nucleosome occupancy while irregular nucleosome positioning was observed across the R1 and R2 templates, with the fibres showing substantial heterogeneity in nucleosome binding between molecules, similar to patterns seen in cells^17^. Notably, analysis of methylation foot printing data revealed that 601 nucleosomes showed pronounced delocalization and a loss of strong positioning when positioned within the R2 template.

### Irregular nucleosome spacing leads to disordered chromatin fibre structures

We previously showed that regularly spaced nucleosome arrays, such as those found in satellite-containing heterochromatin, form compact chromatin structures^8^. In contrast, bulk chromatin is more heterogeneous and interspersed with discontinuities^8,32^. We hypothesised that this heterogeneity arises from irregularly spaced nucleosome arrays that reduce nucleosome stacking. To test this, we characterised the structure of reconstituted chromatin fibres formed using the 601, R1, and R2 sequences. Fibres were assembled either with or without linker histones to produce compacted or unfolded fibres, respectively^33^. To assess chromatin compaction under native-like linker histone levels, we employed an “H5-redistribution” approach that leverages the high mobility of linker histones^34,35^. Nucleosome arrays were spiked into purified chicken erythrocyte (CE) chromatin (-/+H5), which will promote the transfer of H5 from the CE chromatin to the reconstituted arrays until equilibrium is reached, avoiding precipitation caused by LH oversaturation (Extended data 4a-b).

Sucrose sedimentation analysis was used to characterise bulk folding properties of reconstituted chromatin fibres in solution (Extended data 4a-d). The 601 template had the same sedimentation characteristics as CE chromatin, which is known to form canonical “30 nm” like fibres in the presence of linker histones^36^, confirming H5-redistribution (Extended data 4e). In the absence of linker histones these fibres sedimented more slowly, consistent with them having a more open or disrupted structure. Fully reconstituted R1 and R2 fibres with linker histones sedimented more slowly than 601 fibres with a wider peak, consistent with them having a more disrupted and heterogenous structure (Extended data 4f-g)^8^; small angle X-ray scattering data also suggested more heterogeneity (Extended data 5a-b; see Additional Notes).

To characterise structural heterogeneity, transmission electron microscopy (TEM) was used to study the single molecule structure of 601, R1 and R2 chromatin fibres. Biotin-labelled DNA was reconstituted with core and linker histones and separated from donor nucleosome core particles and CE chromatin using streptavidin coated beads (Fig 2a-b). 601 arrays with a canonical 197 bp repeat are known to form compact fibres^37^; accordingly they had a regular fibre structure with a median diameter of 38.5 nm (SD ± 9.1 nm), similar to canonical CE chromatin (Fig 2c, Extended data 6a-b).

**Figure 2.**
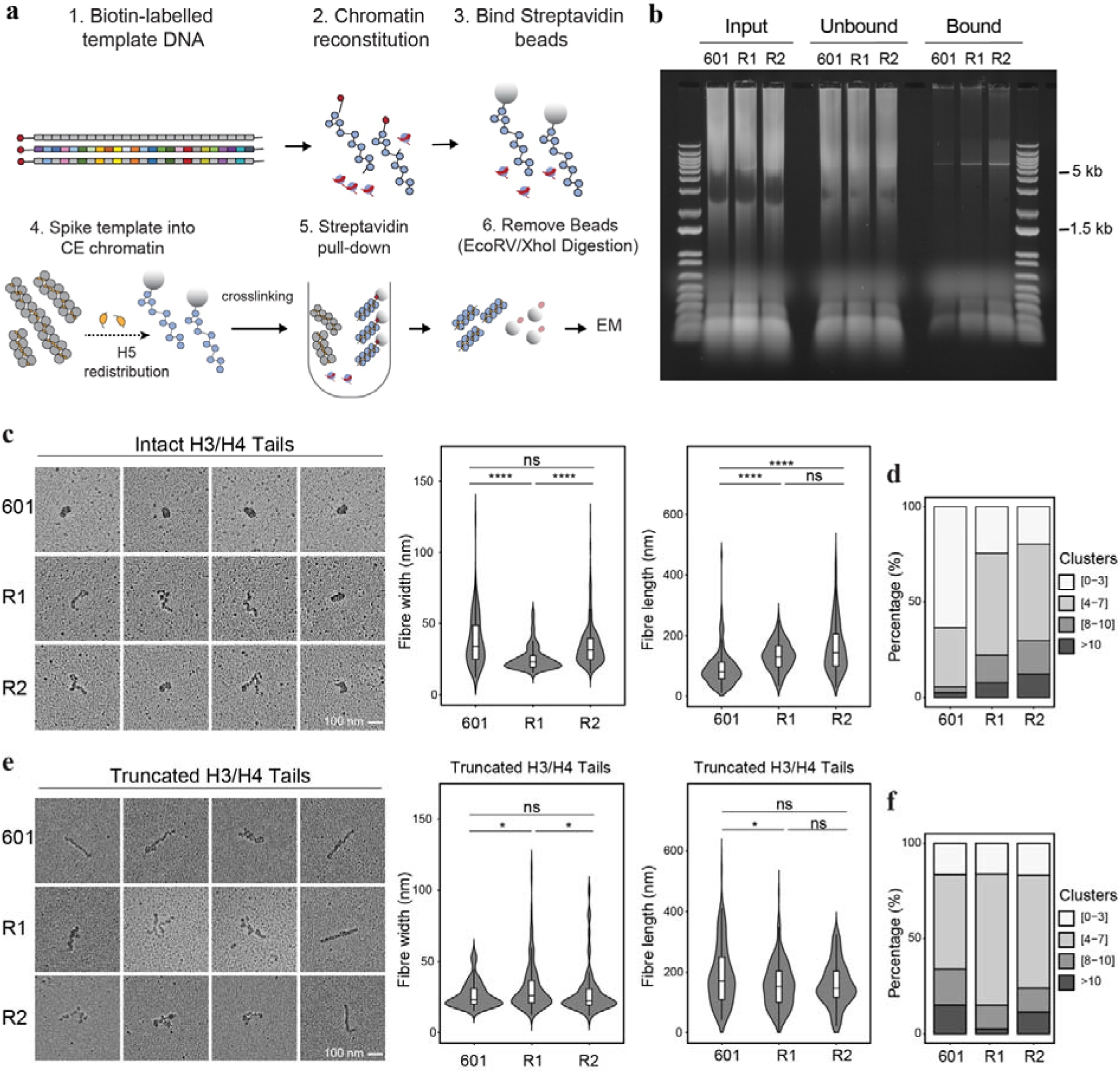
Transmission electron microscopy (TEM) of irregular chromatin fibres. **a,** Schematic showing the preparation of chromatin fibres for TEM. **b,** Agarose gel showing enrichment of reconstituted chromatin for TEM. Input chromatin was bound to beads and folded by H5-redistribution, the unbound fraction was washed off and bound template chromatin (∼5.5 kb) was cleaved off for TEM. **c,** Left: Gallery of representative TEM images of arrays reconstituted with NCPs with intact histone tails and folded by H5-redistribution. Right: Violin plots showing measurements of fibre width and length in arrays reconstituted with wild-type NCPs (601, n = 158; R1, n = 65; R2, n = 316). **d,** Box plots showing frequency of nucleosome clusters within fibres in arrays reconstituted with wild-type NCPs (601, n = 161; R1 n = 213; R2, n = 323). **e,** Left: Panel showing representative TEM images of arrays reconstituted with NCPs with truncated H3/H4 tails, in the presence of linker histone. Right: Violin plots showing fibre width and length in arrays reconstituted with NCPs with truncated H3/H4 tails (601; n = 125; R1, n = 109; R2, n = 79). **f,** Box plots showing frequency of nucleosome clusters within fibres (as in **d**) but reconstituted with NCPs having truncated H3/H4 tails (601, n = 89; R1, n = 287; R2, n = 87). Statistics are the result of a Wilcoxon Test (*, P≤ 0.05; **, P≤ 0.01; ***, P ≤ 0.001; ****, P ≤ 0.0001).

Despite being similarly saturated with histones, R1 and R2 fibres showed extremely heterogenous structures (Fig 2c, Extended data 7), that were confirmed by atomic force microscopy to not be a consequence of fixation conditions (Extended Fig 8a-b). These fibres were less condensed with a median diameter of ∼19-21 nm with linker DNA connecting small nucleosome clusters (Fig 2d), that may correspond to the nucleosome “clutches” observed *in vivo*^11,38^. R1 and R2 irregularly spaced arrays were also more elongated and showed wider variation in local compaction compared to the 601 arrays (Fig 2c, left), which was supported by further analyses of the fibre morphology (Fig 2c, right).

We speculated that differences in fibre structure might be mediated by nucleosome-nucleosome stacking interactions, through histone tails^37,39–41^. To test this, we reconstituted chromatin fibres with NCPs lacking H3/H4 tails (Extended data 9a-c). Loss of H3/H4 tails caused regular 601 fibres to adopt a more extended structure, with reduced diameter and increased fibre length and cluster number (Fig 2e-f), making them more similar to R1 and R2 fibres. These results confirm compaction is mediated by the H4 N terminal tails, presumably interacting with the H2A-H2B acidic patch^42^. In contrast, tail removal had little effect on the number of clusters in R1 and R2 arrays (Fig 2d vs 2f), indicating that these structural motifs were independent of H4/H2A-H2B interactions and may represent an intermediate folding state. Given that 601 derived heterochromatin-like fibres are more compact (Fig 2c), we infer that they rely more strongly on histone tail interactions than euchromatic-like fibres, highlighting how small changes in nucleosome positioning and a weakening of stacking interactions can substantially alter chromatin fibre structure.

### 3D polymer simulations of chromatin fibres reveal how irregular nucleosome spacing leads to disordered structures

TEM showed that euchromatic fibres of the same sequence can adopt very different structures (Fig 2c-f). To gain mechanistic insight into the molecular basis of fibre folding and structural heterogeneity, we developed a coarse-grained quantitative model of chromatin that could be used for molecular dynamics simulations (Fig 3a, Extended data 10). In this model, termed NucHOP (Nucleosome-resolved Higher order Polymer), the DNA and nucleosomes are represented as a series of ‘beads’ which diffuse in solution subject to thermal motion and phenomenological interaction potentials (see Methods for details). Although many nucleosome-resolution computational models have considered fibres formed from nucleosomes with regular nucleosome spacing^43,44^, this new model can accommodate heterogenous nucleosome spacing, obtained from experimental data (Fig 1d). The key model ingredients were a realistic excluded volume for nucleosome core particles and linker DNA, an effective inter-nucleosome interaction potential, DNA elastic properties (bending and twisting), and correct relative orientation of neighbouring nucleosomes based on their spacing. To ensure the simulations were computationally efficient, some simplifications were made: the nucleosomal DNA could not unwrap from the histone core, and the positions along the DNA of the nucleosomes were fixed for the duration of each given simulation. Although linker histones were not explicitly included in the model, their effects were: firstly, by influencing DNA entry-exit angles^45–47^ and secondly by an effective stiffening of the exiting DNA by the linker histone tails^39,48^.

**Figure 3.**
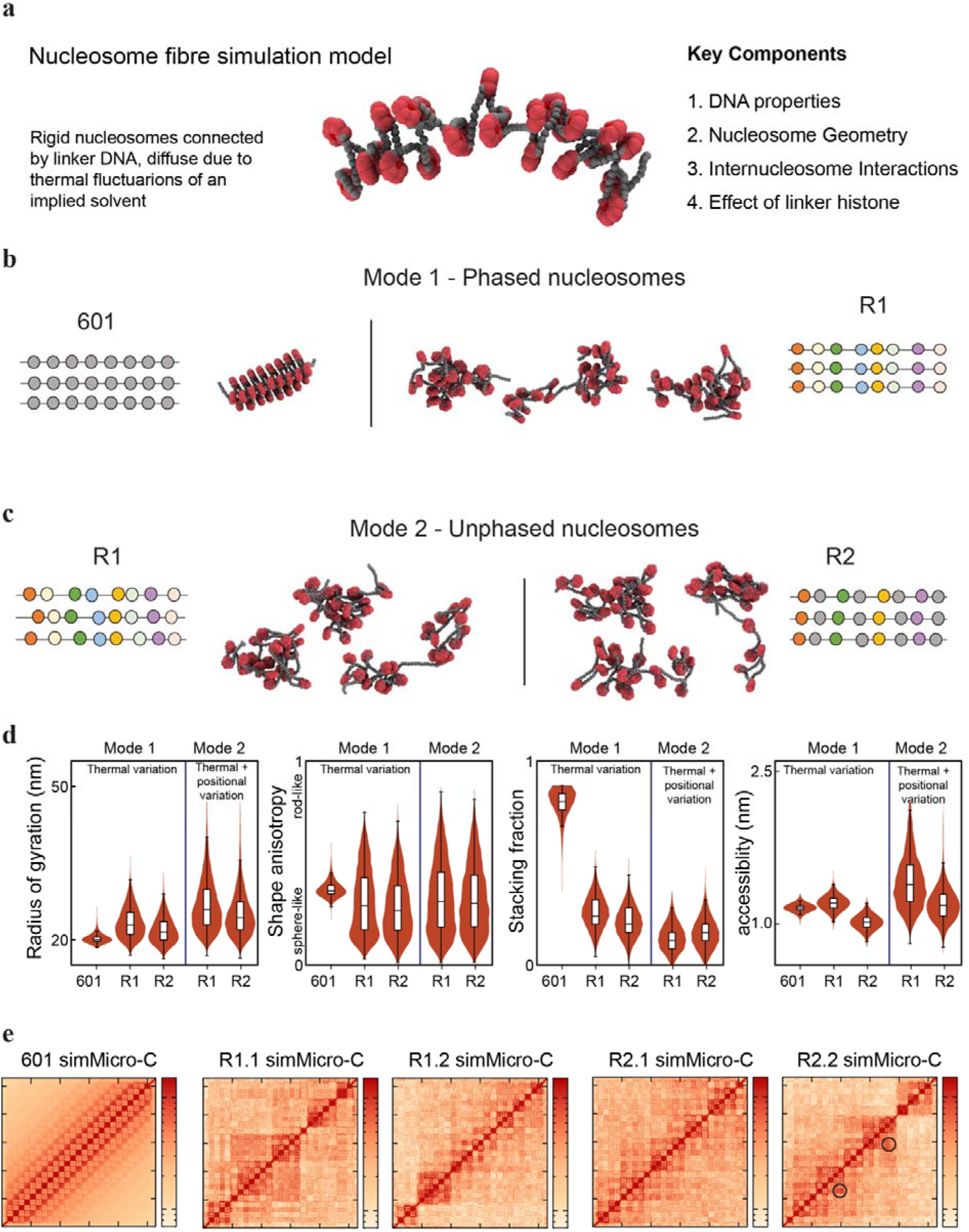
Polymer simulations of irregular nucleosome arrays to quantify fibre heterogeneity. **a,** The 3D structure of nucleosome fibres was modelled using a coarse-grained bead-and-spring polymer model. Four key components are included in the model: DNA biophysics, nucleosome geometry, inter-nucleosome interactions and the effect of linker histones (see Methods for details). Nucleosome positions / linker lengths were obtained from MNase-seq data. **b,** 50 simulations of the same nucleosome fibre were performed for each template, analogous to ‘phased nucleosome arrays’. Each simulation was run for a time equivalent to 36 ms, with 101 configurations (‘snaphots’ in time) extracted from each. **c,** 50 simulations were performed, each for a different R1 or R2 template. These are ‘unphased nucleosome arrays’, which capture fibre-to-fibre nucleosome position variation. **d,** Violin plots showing distributions of the radius of gyration, shape anisotropy, stacking fraction, and accessibility for each simulated chromatin fibre. The radius of gyration and shape anisotropy measure the typical 3D size and shape of the simulated molecule (see Methods for details). The stacking fraction counts the fraction of nucleosomes which are ‘stacked’ together (i.e., involved in close face-to-face nucleosome-nucleosome interactions), and the ‘accessibility’ gives a simple average measure of the space available around DNA segments within a fibre. Each distribution is obtained from a set of 50 simulations (5050 individual snapshots/values) as described in **c** and **d**. The width of the violins is scaled to be the same. Box plots overlaid on the violins show the median and first and third quartiles; whiskers extend to 1.5 times the interquartile range. **e,** Fibre interaction maps (simulated ‘MicroC’) are shown for 601 and examples of unphased R1 and R2 fibres. In each case the contact map is obtained from a single simulation (101 configurations captured over a time equivalent to 36 ms). The horizontal and vertical axes show the bp position along the fibre (same scale on each axis); the darker the colour, the more likely the two points along the fibre are to be found in close spatial proximity during the simulation. Nucleosome^m^ to nucleosome ^m+n^ interactions are depicted by black circles.

Chromatin fibre constructs were modelled in two different ways, phased and unphased. For phased models, we simulated fibres with nucleosome positions which best fit the MNase-seq data for each of the three constructs used. For each construct, 50 independent simulations were performed with a duration equivalent to 36 ms, each with identical nucleosome dyad positions. This enabled an assessment of structural variability over time across copies of the same fibres only affected by thermal fluctuations. Such ‘thermally generated structural variation’ is akin to population variability in active gene promoters within a specific cell type, which have irregular nucleosome spacing but are strongly phased across the cell population^17^.

As observed in the experimental data, 601 fibres have a compact highly folded structure in contrast to phased R1 and R2 arrays (Fig 3b, Extended data 12a), and instead these fibres had pronounced disruptions and clear variations in chromatin folding. To quantify differences in fibre structure we used two intuitive global measures: radius of gyration (a measure of the 3D size) and shape anisotropy (indicates whether the fibre is sphere or rod-like) (Fig. 3d). Two parameters quantified the internal structure variations: the ‘stacked fraction’, and a DNA ‘accessibility’ score. The stacked fraction is the proportion of nucleosome faces which interacted with another nucleosome at any given instant to quantify the degree of liquid crystalline order in a fibre. The accessibility is an estimate of the largest object which could bind the linker DNA without disrupting the local fibre structure (Extended data 13). It was calculated by taking each DNA base-pair of the simulated fibre and measuring the size of the largest sphere which can fit around that base-pair without intersecting with any protein component (nucleosome core) within the simulated configuration. We considered the fibre-level accessibility by taking the mean accessibility across the fibre for a given configuration.

Overall, these measures suggested that thermal motion only modestly affects structural variation in the 601 fibres, while there is much broader structural variability for the irregular R1 and R2 fibres. The stacking fraction measure shows that 601 fibres usually adopt an ordered configuration where most nucleosomes are stacked. Instead, phased R1 and R2 best fit fibres show much lower stacking fractions with fewer than half of the nucleosome faces stacked together. For the 601 fibres, the distribution of fibre-level accessibility is narrow, whereas the standard deviation is more than twice as large for R1 and R2 fibres. Interestingly, the mean value of the fibre-level accessibility for R1 is larger than 601, but smaller for R2 which could be due to differences in linker length composition. (Fig. 3d).

To ask how variable nucleosome positions between molecules affected structure, we then ran another set of simulations using unphased chromatin fibres. For these, a set of 50 simulations for each fibre construct was considered, where in each case the nucleosome dyad positions were unique, but still based on the distribution of nucleosome positions observed in the MNase-seq data (Fig. 1d, Extended data 11). This enabled us to assess the effects of small differences in nucleosome positions between fibres, or the ‘positioning generated structural variation’ (including thermal variations) and best reflects how variations in nucleosome position between different cells might influence chromatin fibre structure^17^. Unphased R1 and R2 fibres were more extensively disrupted, indicating that irregular nucleosome positioning strongly destabilised folding (Fig 3; Extended data 12b-d).

To better characterise nucleosome-nucleosome interactions across fibres we calculated an interaction matrix, analogous, to micro-C (Fig 3e, Extended data 12e). Within a single fibre, e.g. R1 example 1, there were clear nucleosome interaction clusters, or ‘clutches’, that varied between fibres (compare to R1 example 2). Despite R2 having strong 601 nucleosome binding sites these did not dominate interactions, suggesting that single strong nucleosome binding sites do not significantly affect nucleosome-nucleosome stacking and instead the fibres form a series of clutches, similar to R1. For both fibres there are further sub-clutch interactions and, for example R2 example 2, there are symmetrical off-diagonal signals indicative of nucleosome^m^ to nucleosome^m+n^ interactions.

### Irregular nucleosome arrays show dynamic sampling of structures

Simulations can provide dynamic insight into chromatin fibres, which is challenging, or impossible, to obtain experimentally. Individual 601, R1 and R2 fibres were evaluated over time (Movie 1-2), focusing on radius of gyration, shape anisotropy, stacking fraction, and accessibility (Fig 4a, Movie 3). The 601 fibres maintain high levels of stacking interactions, which changed little over time. The other structural parameters also indicated uniform accessibility, radius of gyration and anisotropy. In contrast, R1 and R2 fibres were highly dynamic (Movie 1-3) with the radius of gyration and shape anisotropy fluctuating, coincident with changes in the stacking fraction. Simulation measurements are shown for 7 ms, demonstrating how the fibre samples many different types of structures. In a cellular context these may modulate transcription factor binding site accessibility and provide a platform for chromatin remodelling machines to function within an already dynamic landscape.

**Figure 4.**
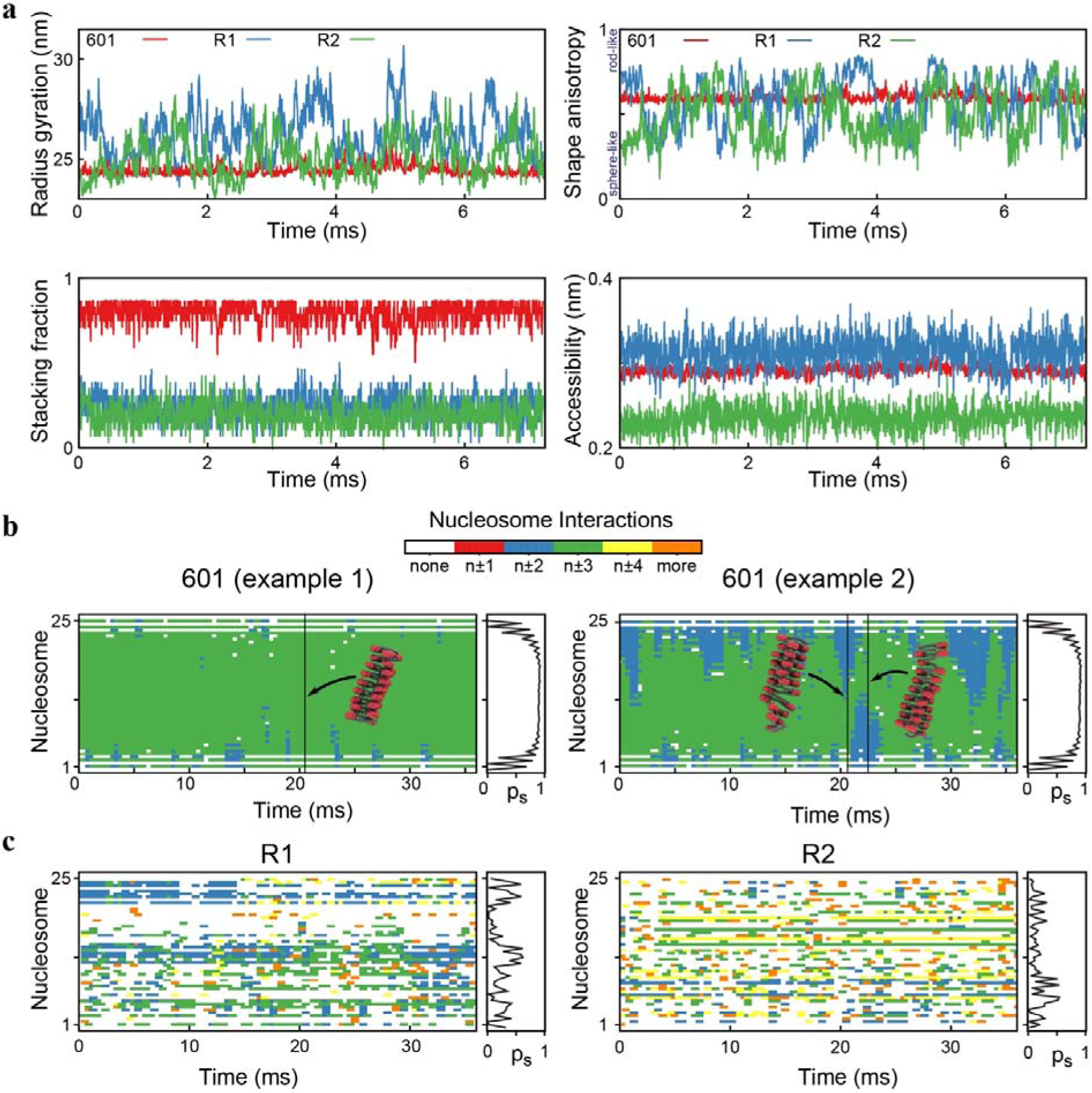
Nucleosome array dynamics revealed by polymer simulations. **a,** Plots showing how different structural parameters (radius gyration, shape anisotropy, stacking fraction and accessibility) vary with time as measured in three example simulations of phased 601 (red), R1 (blue) and R2 (green) fibres. The simulations were run for the equivalent of 7.5 ms, with the configuration saved every 0.0018 ms. **b,** Plots of 601 fibres showing how the nucleosome stacking interactions change with time within individual fibres. A face of a nucleosome is ‘stacked’ if it is within a threshold separation of another nucleosome face. At each time point (horizontal axis), the stacking partner of a given face (vertical axis) is indicated by the colour. Insets show fibre structures at these timepoints. The probability, p_s_, that a nucleosome is found interacting with its most common partner is shown to the right of each plot. **c**, As for (**b**), but instead showing plots for R1 and R2 fibres.

Above we demonstrated that nucleosome-nucleosome stacking interactions were a major determinant of chromatin compaction (Fig 3d), mediated by the linker histone tails (Fig 2e). To better explore this, stacking interactions were analysed over time during chromatin fibre simulations (Fig 4b). For the 601 fibres most interactions were n±3, giving rise to regular arrays (Fig 4b, inset), whilst transitions to n±2, introduced small disruptions in the fibre (see example 2). In contrast nucleosome-nucleosome interactions in the R1 and R2 fibres varied considerably over time (Fig 4c), fluctuating between interactions with n±1,2,3,4 and more, with interactions away from n±3 giving rise to deviations in chromatin fibre folding creating points of accessibility. This was even more pronounced in the unphased fibres compared to phased fibres (Extended data 14a-d), as seen in the p_s_ curve (probability that a nucleosome is found interacting with its most common partner). This indicated that there were very few preferential interactions, suggesting there will be significant structural variability between different copies of the same loci between cells.

### Disordered chromatin fibres are deformable

Experiments (Fig 2) and simulations (Fig 3) showed that irregular nucleosome spacing leads to disordered chromatin fibre structures. To investigate how this disorder shapes the mechanical properties and unfolding pathways of the fibres, single-molecule force spectroscopy was performed by multiplexed magnetic tweezers (MMT). This setup applies controlled forces to many individual chromatin fibres in parallel, allowing detection of transient structural changes and resolution of molecule-to-molecule variability within heterogenous samples^22,49,50^. There are two regimes observable for chromatin folding: at low forces fibres behave like springs, reflecting an elastic extension followed by nucleosome unstacking and partial DNA unwrapping from the histone octamer^50^. At higher forces, histones are fully unwrapped leading to stepwise extension of the fibres^49,50^. The low force regime is most relevant for investigating disruptions in chromatin folding, where chromatin-associated machines might exert force that destabilise higher-order folding^51^.

Chromatin fibres were reconstituted with NCPs on template DNAs end-labelled with biotin and DIG (Fig 5a-b). Fibres were tethered between a glass slide surface and superparamagnetic beads, and then CE chromatin (+H5) was flushed in to facilitate H5-redistribution. Force-extension curves (Fig 5c, Extended data 15a) for 601 fibres showed that they were able to resist force below 2 pN before undergoing a significant change in extension at higher forces. In contrast, at low forces (< 2 pN) R1 and R2 fibres were readily extended, indicating that the fibres are more fragile, presumably due to reduced nucleosome–nucleosome interactions.

**Figure 5.**
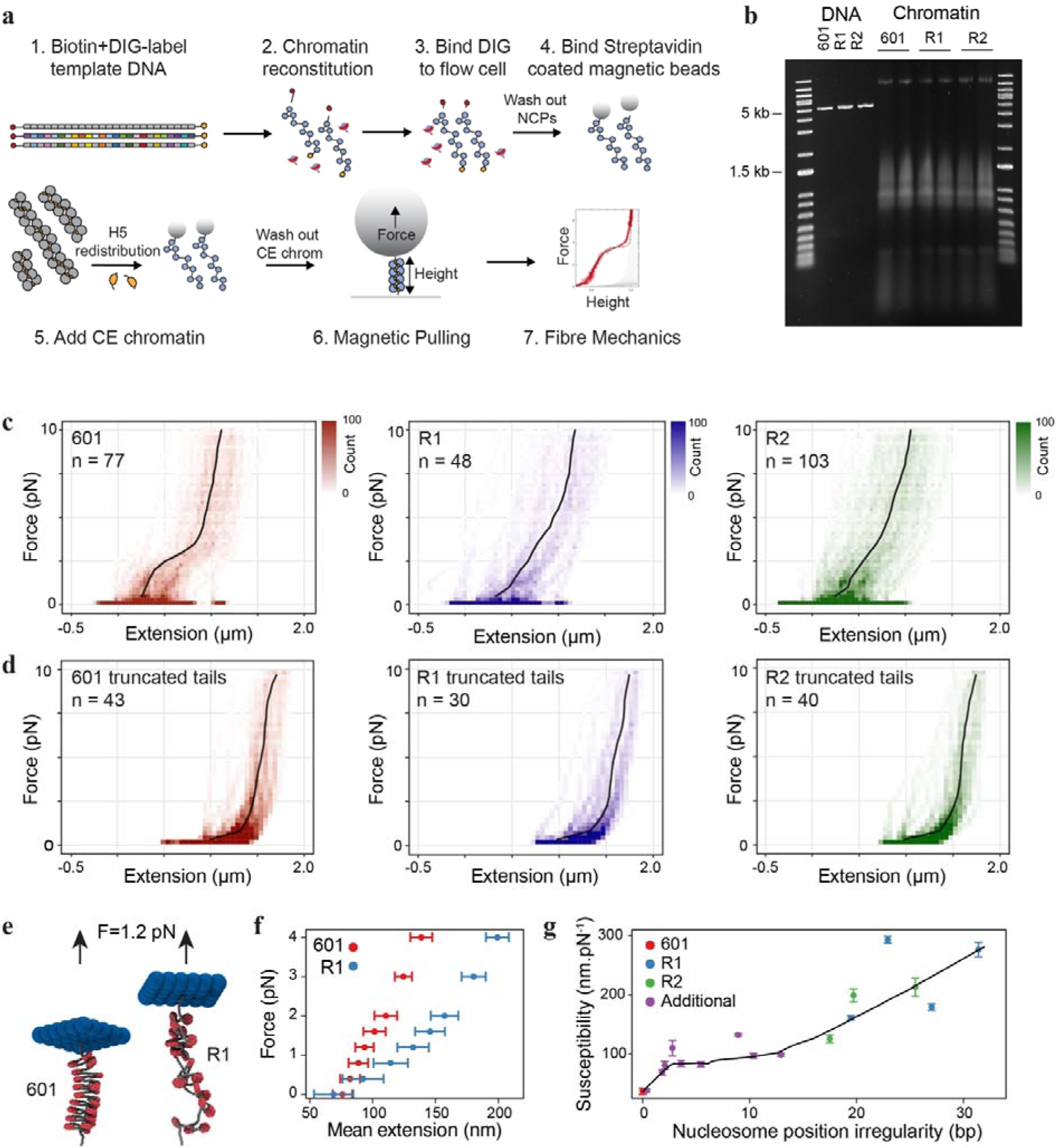
Irregular nucleosome arrays are fragile under tension. **a,** Diagram showing the reconstitution of chromatin fibres on biotin and DIG-labelled DNA templates. After reconstitution, fibres were passed through a flow cell coated in anti-DIG antibodies, superparamagnetic beads and CE chromatin (+H5) were then flowed through. Individual molecules were extended using a magnetic field gradient, and the tether height measured by microscopic imaging of the bead spatial coordinates in real time. **b,** Electrophoretic mobility shift assay (EMSA) showing retarded migration of 601, R1 and R2 biotin and DIG-labelled templates after chromatin reconstitution by histone-transfer. **c,** Heat-maps depicting meta-analysis of force-extension curves for reconstituted chromatin fibres folded with linker histones by H5-redistribution. Black line corresponds to median data. **d,** Same as (**c**), except fibres were reconstituted with NCPs with truncated H3/H4 tails. **e,** Force-extension behaviour of simulated fibres probed using an arrangement where the fibre is fixed at one end, and the other end is attached to an array of beads (blue). A constant force is applied to the array of beads, and the fibre end-to-end distance is measured over time (see Methods for details). Snapshots of 601 and R1 fibres are shown for an applied force of 1.2 pN. **f,** Force-extension curves generated from simulated pulling of 601 (red) and R1 (blue) fibres. Each point is an average (n=4200 over 3 independent simulations), horizontal bars show standard deviation and indicate variation due to thermal fluctuations. **g,** Plot showing susceptibility – a measure of the extent to which a fibre responds to applied force – as a function of the nucleosome position irregularity (mean squared deviation of the linker lengths away from 50 bp, see Methods). For a set of simulated fibres including 601, R1 and R2 unphased fibres, and ‘additional’ fibres with linker lengths generated randomly, see Extended Data Figure 15. Black line corresponds to a LOWESS fit (bandwidth 0.45), vertical bars show standard error of the mean.

To investigate how nucleosome stacking interactions directly affect fibre fragility, chromatin fibres were reconstituted with NCPs lacking H3/H4 tails (Extended data 9). Force-extension curves for these fibres exhibited a remarkably different shape (Fig 5d, Extended data 15b). Firstly, in the absence of H3/H4 tails 601, R1 and R2 fibres were more extended, as they were not fully folded, as observed before by TEM (Fig 2e). Second, 601 fibres became extremely fragile and showed little resistance to force, similar to R1 and R2 fibres, and more typical of how naked DNA behaves^49^. R1 and R2 fibres, even in the presence of histone tails have few nucleosome-nucleosome interactions (Fig 4b) – indicating that nucleosome interactions are crucial for stabilising intra-fibre structure, leading to fibre stability.

To further understand the relationship between nucleosome stacking and susceptibility to force, we simulated the effect of pulling on modelled chromatin fibres (Fig 5e). 601 fibres were resistant to a pulling force of 1.2 pN whilst R1 fibres were readily extended (Movie 4).

Simulation force extension curves (Fig 5f) indicate that, as in the experiments, fibres with extensive nucleosome-nucleosome interactions were refractory to force. To further understand this relationship, we calculated the nucleosome position irregularity, the mean squared deviation of the linker lengths away from 50 bp, for a series of fibres (Extended data 11, see Methods), and modelled the susceptibility to pulling. Fibres with little nucleosome position irregularity were highly resistant to pulling forces, compared to more disrupted R1 and R2 fibres (Fig 5g, Extended data 15 c-d).

### Nucleosome positioning irregularity establishes distinct structural phase transitions

Molecular tweezer experiments indicated that nucleosome position irregularity correlated with a fibre’s susceptibility to deformation (Fig 5g). To understand the molecular basis for these biophysical characteristics, we examined how stacking energy (calculated as mean stacked fraction) varies with nucleosome position irregularity (Fig 6a). We used additional simulated fibres to fill the gap between 601 fibres, which have zero irregularity, and the R1 and R2 fibres which exhibit irregularities greater than 10 (Extended data 11). 601 fibres display high stacking energy which drops sharply with only 2-3 bp of irregularity, reaching a pronounced plateau observed for R1 and R2 fibres, suggesting that even minimal irregularity is sufficient to disrupt stacking and increase fibre deformability (Fig 5c). We then analysed fibre accessibility (Extended data 13) for different levels of nucleosome irregularity. Fibre accessibility was low for 601 fibres with a functional transition at 18 bp of irregularity, indicating that more pronounced global disruptions in structure are required to facilitate transcription factor binding (Fig 6b). Concomitantly, a transition in fibre deformability was observed at this point (Fig 5c).

**Figure 6.**
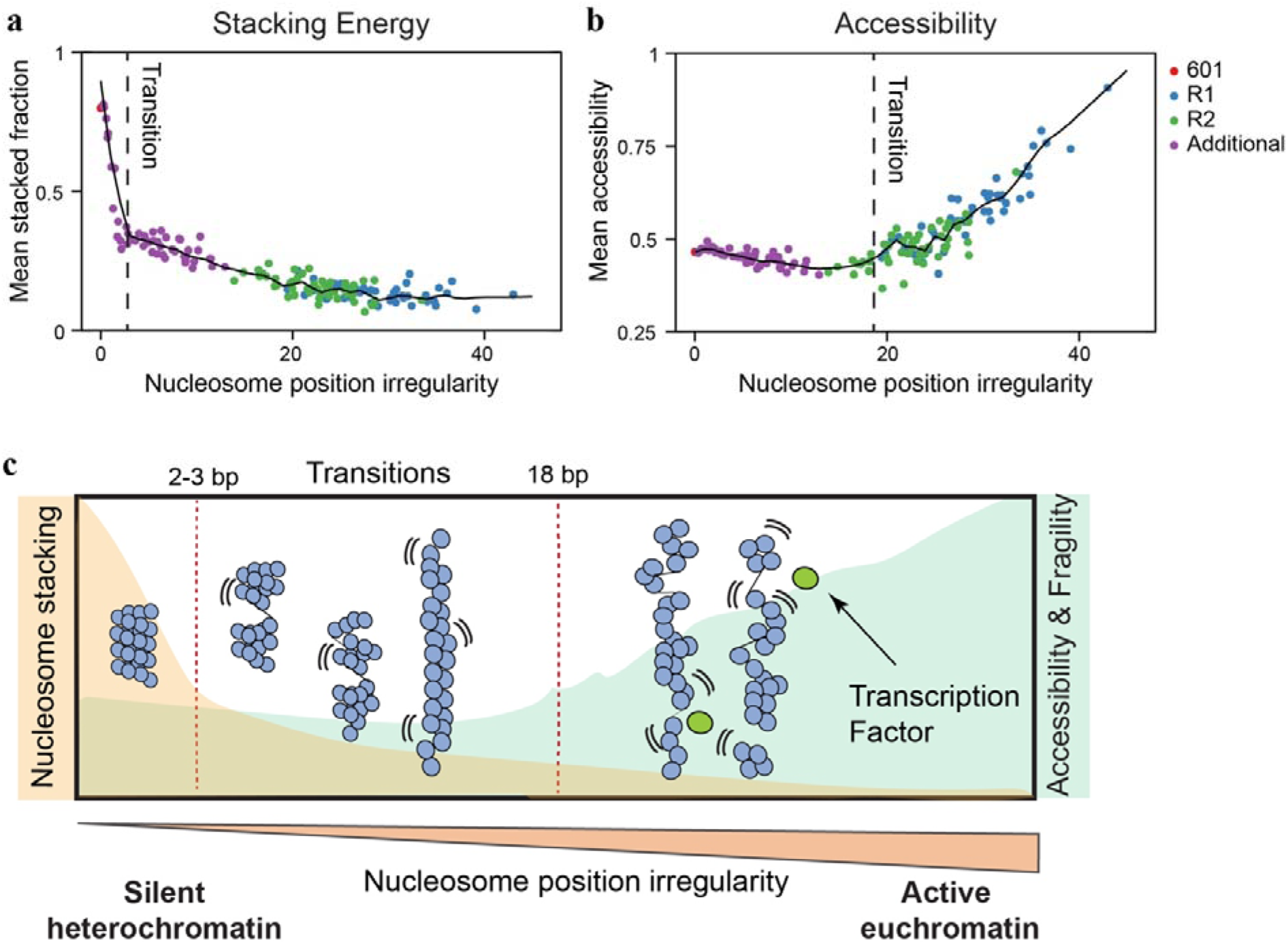
Nucleosome positioning irregularity induces a chromatin phase transition. **a,** Each point is obtained from a single simulation (stacking fraction is taken from 101 snapshot configurations from a simulation duration equivalent to 36 ms). Black line is a LOWESS fit (bandwidth 0.1). **b,** Graph showing mean accessibility as a function of nucleosome position irregularity. Black line is a LOWESS fit (bandwidth 0.1).

## Discussion

This study describes a mechanistic framework for how two distinct transitions in chromatin organization balance genome protection with chromatin accessibility (Fig. 6c). We show that minor irregularity (2–3 bp) in nucleosome positioning along the DNA drives a transition from a para-crystalline phase, in which chromatin fibres fold in an orderly manner, to a disordered, liquid-like phase marked by chromatin decompaction and loss of nucleosome–nucleosome stacking interactions. Importantly, a second functional transition that establishes global fibre accessibility and enhanced fibre fragility requires much greater nucleosome array irregularity (∼18 bp). This suggests that chromatin fibres can accommodate local structural disruption while remaining inaccessible to regulatory factors. Furthermore, this transition is accompanied by a marked increase in structural heterogeneity (Fig 2c) helping to reconcile discrepancies between classical ordered models derived from repetitive sequences^37,42,49,52^ and *in situ* evidence for heterogeneous chromatin fibres in living cells^10,11^.

These structural insights also have direct implications for the sequence-to-expression conundrum: the challenge of understanding how DNA sequence gives rise to quantitative gene expression patterns, given that sequence encodes both biochemical information (e.g. transcription factor binding sites) and biophysical information (nucleosome positioning, chromatin structure)^53^. We suggest that DNA sequence tunes nucleosome positions to adjust the structural state of chromatin, thereby regulating processes such as transcription factor binding^54,55^. Accordingly, we find that euchromatic sequences lie close to the transition between the low- and high-accessibility phases (Fig 6c), where fluctuations are expected to peak near a critical point^56^. This provides a biophysical rationale for why small positional shifts can have disproportionate effects on accessibility and transcription potential.

Structural differences arising from different nucleosome positioning patterns (for example, for the same gene in different cell types) could contribute to cell-type specific gene expression patterns^17^. Within active genes of the same cell type, stochastic fluctuations in nucleosome spacing may also contribute to variation in expression levels, augmenting transcriptional noise^17,57,58^. Current sequence-to-expression (S2E) models often overlook such inherent variability^59^, possibly due to variations in chromatin structure not being well considered. We suggest that future S2E models will account for nucleosome positioning and concomitantly chromatin accessibility landscapes to better predict transcription factor binding and transcriptional output^57,58,60,61^. The reconstitution approaches developed here can also be used to reconstitute other genomic regions, and the NucHoP chromatin model could be extended to model regions up to 20 kb in length, enabling larger segments of the genome to be studied in future.

Single-molecule stretching experiments revealed that irregular nucleosome spacing increased fibre fragility under tension (Fig 5). Together with the enhanced structural variability of non-repetitive arrays, this feature may facilitate fast structural sampling and dynamic remodelling of local chromatin (Fig 4), important for nuclear processes such as transcription. Indeed, the force range used in magnetic tweezer experiments lies within the range of forces applied by molecular motors such as RNA polymerase II and the RSC remodelling complex^51^. Therefore, the observed fragility may promote rapid changes in chromatin structure mediated by the action of the transcription machinery or of chromatin remodelling complexes.

We suggest there is a need to shift towards chromatin models that account for irregular nucleosome positioning, as only these lead to the inherent disrupted, fragile and highly heterogeneous chromatin structure which are relevant *in vivo*. Irregular nucleosome spacing shapes not only fibre compaction but also temporal variability and susceptibility to local perturbation—highlighting the importance of considering chromatin dynamics, not just static structure, to fully understand the physical basis of gene regulation.

## Acknowledgements.

We thank Duncan Sproul for useful comments on the manuscript. We also thank members of the Gilbert and Marenduzzo groups for ongoing interesting discussions on chromatin architecture. This work was supported by the European Research Council (CoG 648050 THREEDCELLPHYSICS) to D.M., N.G. was supported by the UK Medical Research Council (MC_UU_00035/6). N.G and D.M. are Wellcome Investigators (223097/Z/21).

## Author contributions

NG and JA conceived the project; NG, DM, J van N and CB supervised the project. WA, G-JK, HW, AK, JD, WV undertook lab-based experiments. MT, MC, CB, did the computational simulations. All authors wrote the manuscript.

## Additional notes

### Small angle X-ray Scattering

Small angle X-ray scattering (SAXS) was used to examine the folding envelope of the chromatin fibres. As SAXS requires highly pure material for analysis, it was incompatible with using the H5-redistribution method, instead fibres were folded by the direct addition of purified CE H5 to a 1:1 H5:nucleosome ratio. In the absence of linker H5, 601 fibres showed a peak corresponding to a periodicity of around 19.5 nm (q = 0.05) and a small shoulder at around 12 nm (q = 0.083) (Extended data 5). These peaks have been previously observed in 601 arrays^62^, and likely reflect the periodicity of the nucleosome face and the linker DNA connecting the nucleosomes, as the nucleosome width is ∼11 nm, and the linker length in these arrays is ∼50 bp (∼17 nm). In the R1 and R2 fibres, this peak is less prominent, resulting in a broad peak centring around 17-19 nm in the R1 fibres and 18-22 nm in R2 fibres reflecting the variability in linker lengths within BLG sites. In the presence of H5, 601 fibres showed a gradual reduction in the 12 and 20 nm peaks, and the appearance of a shoulder at a larger periodicity of ∼60-66 nm. This is consistent with compaction into a higher-order structure, although interestingly a 30 nm peak was not observed, as reported previously^12^. Similarly, the R1 and R2 fibres showed a shift in the direction of a larger periodicity in the presence of H5. The R2 fibres appear to be more compacted, with a periodicity in the range of 23-40 nm, compared to approximately 17.5-27.5 nm in R1 fibres. Importantly, R1/R2 arrays maintained a broad peak distribution indicating a larger heterogeneity in 3D structure within the population, and there was a pronounced difference between R1/R2 and 601 fibres, suggesting there were large differences in secondary structure.

### Polymer simulations

While developing our model, we found that the specific folded structure of the 601 fibres depended strongly on the model parameters. For example, these fibres can fold into both 2-start and 3-start solenoid type structures, which is most predominant depends on the precise entry-exit angle of the DNA at the nucleosome dyad. Though we have parametrised the model using experimental data, the limitations of the experimental parameter measurement and of the simplified course-grained simulation approach mean that the model may not be able to predict precisely which folded structures are the most likely to form. This will depend on details such as the linker histone, core histone tails, and the presence of mono- and/or di-valent salts, which are not fully captured by the model. Nevertheless, such details are unlikely to change the behaviour of the model in terms of differences between the regular and irregular fibres.

For simulated single-molecule force spectroscopy the simulation geometry is different from the experiments. Consequently, the magnitudes of force and extension cannot be directly compared; furthermore, in the simulations DNA cannot ‘unwrap’ from the nucleosomes, and so they can only access the regime where nucleosome unstacking is the dominant source of increasing extension. It is important to emphasise that this is the most relevant structural change at a low force regime.

## Methods

### Nucleosome positioning array synthesis

A pUC18 plasmid containing 26 repeats of the 601 sequence (26 x 197 bp) was kindly provided by Daniela Rhodes^37^. The number of repeats within the plasmid is inherently unstable due to the repetitive nature of the template; and individual clones grown at 37°C following transfection into DH5α cells contained between 19 and 26 repeats of the “601” sequence, however, no plasmids containing 25 repeats could be identified. It is possible that a single copy of the 197 bp repeat is never deleted due to constraints in the bending of the DNA required for the recombination event leading to deletion. A plasmid containing 25 copies of the 601 sequence (*601*) and one with 24 copies of the “601” sequence and a central low affinity nucleosome (*601-LA*) were constructed as follows: A plasmid containing 24 repeats of the 197 bp “601” site was partially digested with *Ava*I, which cuts between each of the “601” nucleosome positioning sites. Linearised sequences containing 24 “601” sites were isolated by gel extraction from NuSieve GTG Agarose (Lonza) with an E.N.Z.A. gel extraction kit (Omega), and the 5’ base was dephosphorylated by antarctic phosphatase (NEB). A 3’ phosphorylated 26 bp DNA fragment containing two *Bsg*I restriction sites was ligated to the fragment and used to recircularise the plasmid. Plasmids were transformed into Stbl2 competent cells (Thermo Fisher), clones were grown on agar plates, colonies were picked and DNA prepared using a miniprep kit, (Qiagen). *EcoR*V and *Bsg*I digestion were used to identify clones where the insert appeared in the centre of the template, with 12 “601” sequences on each side. These plasmids were cut with *Bsg*I, and linear dsDNA (GeneArt Strings by Thermo Fisher) containing either “601” or a low affinity (LA) positioning sequence were cloned into the plasmid, such that the 197 base pair repeat was maintained, *Bsg*I restriction sites were lost, *Ava*I restriction sites between nucleosome positioning sites were reformed, and the inserted “601” sequence was indistinguishable from the other repeats. For the R1 and R2 templates, BLG positioning sites were identified from MNase-seq maps of the full length BLG gene^26^, and R1 and R2 plasmids were synthesized by GeneScript in a pUC57 vector. Plasmids were transformed into Stbl2 cells and grown at 30°C to avoid recombination of repetitive sequences. For the excision of the template DNA, *EcoR*V-HF (for 601 and 601-LA) or *XhoI*I (for R1 and R2) restriction enzymes were used.

### Large-scale template DNA purification

To remove the vector backbone from the 601, 601-LA, R1, and R2 templates, approximately 1 mg of 601 and 601-LA plasmid was digested with *Eco*RV*-*HF, *ApaL*I, *BspH*I, and *Eci*I, while R1 and R2 plasmids were digested with *Xho*I*, ApaL*I, *BspH*I, and *Eci*I. Samples were digested overnight at 37°C shaking at 650 rpm to fragment the vector backbone into fragments of less than 800 bp. The digestion was checked for completeness the next day on a 1% (w/v) agarose gel. The sample was purified by phenol:chloroform extraction and ethanol precipitation and resuspended in TE+50mM NaCl buffer (10 mM Tris-HCL pH 7.6, 1 mM EDTA, 50 mM NaCl). The insert was purified from the vector by fractionation with size-exclusion chromatography using a Sephacryl S-1000 column. 40 fractions were collected every 2.5 mins and run on a 1 % (w/v) agarose gel. Fractions containing the insert alone were pooled and purified by ethanol precipitation and resuspended in dd.H_2_O or TE buffer. The concentration of the sample was determined by the Qubit Fluorometer and DNA was stored in −20°C.

### Nuclei preparation

Nuclei were purified from chicken erythrocytes as described previously^63^. All steps were undertaken at 4°C. Briefly, 100 ml of chicken blood was collected and diluted to 360 ml in PBS+Heparin buffer (0.5 mM EGTA, 0.2 mM PMSF, 25 U/ml Heparin in PBS). Blood was centrifuged at 2000 g for 5 mins at 4°C. The supernatant and buffy coat layer were aspirated and red cell pellet was washed and resuspended with 180 ml PBS+Heparin buffer and centrifuged as above. The supernatant was removed and pellet was washed and spun as above. The supernatant was aspirated and the pellet was resuspended in a minimum volume of Buffer A (0.25 M Sucrose, 6 mM MgCl_2_, 50 mM Tris-HCl pH 7.5, 0.5 mM EGTA, 0.2 mM PMSF). The resuspended sample was added dropwise to a beaker containing 800 ml Buffer A+1 % (v/v) Triton X-100 and stirred gently for 18 mins. The sample was split into Sorval bottles and spun as above. The supernatant was aspirated and the nuclei pellet was washed with 100 ml Buffer A. An aliquot was digested with DNase I and the absorbance at 260 nm was measured. Nuclei were diluted to 360 ml in Buffer A then spun at 2000 g for 3 mins. The supernatant was aspirated and the pellet was resuspended in 100 ml MNase digestion buffer (0.2 M Sucrose, 1 mM CaCl_2_, 5 mM Tris, 80 mM NaCl, 0.1 mM PMSF). To lyse the nuclei and remove the nuclear membranes, nuclei were partially digested with 30 U/ml MNase (Worthington) for 7 mins at room temperature with gentle rotation. The reaction was quenched by adding EDTA to a 2.5 mM final concentration and chromatin was stored on ice. 4 ml of this material was used for CE chromatin preparation while the rest was used for the NCP preparation.

### Chicken erythrocyte chromatin preparation

Chromatin was released from nuclei by dialysing overnight into TEP80 buffer (10 mM Tris-HCl pH 8.0, 80 mM NaCl, 0.2 mM EDTA, 0.2 mM PMSF) at 4 °C in using 15.9 mm Visking dialysis tubing (12-14 kDa MWCO). Dialysed chromatin was centrifuged for 2 mins at 5000 rpm to remove nuclear debris, and the supernatant was loaded on 6-40% (w/v) linear sucrose gradients prepared in TEP80 and spun for 2.5 hours at 41,000 rpm at 4°C. Fractions were collected every 0.5 mins and then aliquots were analysed on a 0.7% (w/v) agarose gel and fractions containing chromatin approximately 2.5 to 15 kb in size were pooled, and stored at −20°C. For H5-redistribution experiments chromatin was prepared by thawing on ice and then dialysing to TEP80 buffer overnight at 4°C using 15.9 mm Visking dialysis tubing (12-14 kDa MWCO). Chromatin was then concentrated to approximately 1 mg/ml using Amicon® Ultra-4 Centrifugal Filter Units (Sigma-Aldrich) and stored on ice. For experiments with stripped linker histone, chromatin was stripped of linker histone H5 using two cycles of DNA-cellulose chromatography.

### Purification of chicken erythrocyte nucleosome core particles

The salt concentration of nuclei (prepared as above) was adjusted to 650 mM NaCl by gradual addition of 5 M NaCl. Samples were then diluted to 60 ml using TEP650 buffer (5 mM Tris-HCl pH 7.5, 0.1 mM EDTA, 0.65 M NaCl, 0.2 mM PMSF). To separate the H1 and H5 from oligonucleosomes, 6 × 10 ml aliquots were pelleted through a 20 ml step gradient of 15% (w/v) sucrose onto a 2 ml cushion of 45% (w/v) sucrose prepared in TEP650 buffer, and gradients were centrifuged in a Beckman Coulter SW28 rotor for 15 hrs at 25,000 rpm at 4°C. Four fractions were collected from each gradient with a volume of 10 ml, 10 ml, 7.5 ml, and 4.5 ml respectively, and the OD260 was measured for each fraction. Fraction 4 from all gradients was pooled and dialysed to TEP80 buffer. Samples were digested with 10 U/ml of MNase (Worthington) for 30 mins at 37°C to yield mostly mononucleosomes, and the reaction was quenched by addition of EDTA to 5 mM final concentration. Core particles were concentrated on an Amicon® Ultra-15 Centrifugal Filter Units (Sigma-Aldrich), and DNA and protein quality were assessed by agarose gels and SDS-PAGE.

### Chromatin reconstitution by histone-transfer

Template DNA was incubated with purified chicken erythrocyte nucleosome core particles (NCPs) (H1-depleted) at a 1:1, 5:1, 10:1, 20:1, 50:1 NCP:template mass ratio in TEP1.6 buffer (10 mM Tris-HCl pH 7.6, 0.2 mM EDTA, 0.2 mM PMSF, 1.6 M NaCl) and 0.25 μg/ml BSA for 20 min at 37°C. Samples were then transferred to Slide-A-Lyzer mini-dialysis caps (10K MWCO) (Life Technologies) and dialysed to TEP600 (10 mM Tris-HCl pH 7.6, 0.2 mM EDTA, 0.2 mM PMSF, 600 mM NaCl) buffer for 3-5 hour and then dialysed overnight to TEP80 or TEP10 buffer (10 mM Tris pH 7.6, 0.2 mM EDTA, 0.2 mM PMSF, 80/10 mM NaCl) at 4°C to produce folded or unfolded arrays, respectively. Typically 5 μg of template DNA was reconstituted in each dialysis cap in a volume of 100 μl. Sample concentration was measured using a Nanodrop against the blank buffer. For experiments requiring pure template chromatin, donor NCPs were removed by separation on a CL-4B chromatography column. Samples were loaded on the column in 70–150 μl volume and 30 fractions were collected every 13 seconds with a peristaltic pump at 4°C. Fractions were analysed on a 1% (w/v) agarose gel in 0.5 x TBE buffer to check for NCP separation, and pure template fractions were pooled. Pooled fractions were concentrated using the Amicon. Ultra-0.5 Centrifugal Filter Units (Sigma-Aldrich) following manufactures instructions.

### Electrophoretic mobility-shift assay (EMSA)

0.7% (w/v) agarose gels were prepared without ethidium bromide (EtBr) in 0.25× TBE buffer and left to polymerise. Samples were loaded into wells in Orange Loading Dye (10% sucrose, 0.25×TBE, OrangeG) and run for 3-3.5 hr at 100 V at 4^°^C. Gels were stained with 0.5 μg/ml EtBr and imaged with a camera on a UV light box (Syngene) or scanned using a Fuji FLA-5100 laser scanner. The 1 kb Plus ladder (Invitrogen) was used as a marker unless otherwise mentioned.

### Core histone saturation assay

Reconstituted chromatin was digested with *Bst*XI (NEB) (1 unit/ μg) at 37°C. Aliquots were taken at different time-points and the reaction was stopped by addition of purple loading dye (containing 0.1% SDS) (NEB). Samples were run on a 1% agarose gel and visualized with a Fujifilm FLA-5100 laser scanner. Densitometric analysis was performed using the ImageJ software.

### Transmission electron microscopy (TEM) of unfolded fibres

For imaging of unfolded fibres, chromatin was reconstituted at a 20:1 NCP:template ratio in TEAP1 buffer (10 mM TEA pH 8.0, 1 mM NaCl, 0.2mM EDTA, 0.2 mM PMSF). Chromatin was crosslinked with 0.1% (v/v) glutaraldehyde for 20 minutes at 4°C with gentle rotation. To quench glutaraldehyde and remove core particles from the template, size-exclusion chromatography was performed using a Sepharose CL-4B column in TEP1 buffer (10 mM Tris pH 7.6, 1 mM NaCl, 0.2 mM EDTA, 0.2 mM PMSF). Pure template fractions were pooled and concentrated, and chromatin concentration was adjusted to approximately 2 ng/μl for use in electron microscopy. Chromatin was coated in benzalkonium chloride (BAC) by incubating in 2×10^−4^ % (v/v) BAC for 1 hour at room temperature, and 5 μl of sample was adsorbed onto carbon/formvar 200 mesh grids (TAAB technologies) for 5 minutes then washed with ddH2O two times. The grids were blotted gently on filter paper then placed on a 50 μl drop of 90% (v/v) ethanol for 3 seconds and blotted again. Grids were transferred to a filter paper and left to dry. To enhance contrast a 3 nm layer of platinum was evaporated onto the grids using the Leica EM ACE600 High Vacuum Sputter Coater at a 7° angle. Grids were imaged on the JEOL JEM-1400Plus Transmission Electron Microscope at 15,000-25,000× magnification.

### Mass spectrometry

Protein samples for mass spectrometry analysis were obtained by acid extraction, precipitation and resuspended in water. Mass spectrometry analysis was done on the Bruker Daltonics 12T SolariX Fourier Transform Ion Cyclotron Resonance Mass Spectrometer (FT-ICR MS) using a C18 Peaches LC column. The clipping sites were estimated based on known cleavage sites and the detected molecular mass of the protein.

### MNase-seq

R1 and R2 template DNA was reconstituted with 20:1 NCP:template ratio in TEP10 buffer (10 mM Tris pH 7.6, 0.2 mM EDTA, 0.2 mM PMSF, 10 mM NaCl). NCPs were removed by running on the CL-4B chromatography column as above, and template fractions were pooled and concentrated on Amicon® Ultra-4 centrifugal filter units. Chromatin was digested with 0.15 U/μl of MNase (NEB) in the presence of 2 mM CaCl_2_ for 30 and 60 mins at 25°C and then for 3 minutes at 37°C. The reaction was stopped by the addition of 2× genomic lysis buffer (300 mM NaCl, 20 mM EDTA, 1 % (w/v) SDS). Proteins were digested using proteinase K for 1.5 h at 55°C and DNA was extracted by phenol:chloroform extraction followed by ethanol precipitation. DNA was fractionated on a 1.6 % agarose gel in 1× TAE buffer and imaged on the MS Safe Blue Electrophoresis System. Bands representing mononucleosome fragments were excised and gel purified using the QIAGEN Gel Purification kit (QIAGEN). Mononucleosome fragments were end-repaired and sequencing libraries were prepared using the NEBNext® Ultra^TM^ II DNA Library Prep Kit (Illumina) following the manufacturer’s instructions. Libraries were sized and quality controlled on the D1000 Tapestation (Agilent), and 150 bp paired-end sequencing was performed on a NovaSeq 6000 sequencing platform (Novogene Cambridge).

Paired end sequencing reads were checked for quality using FastQC, FastP and FastScreen, and mapped to the R1 and R2 template sequence using BWA and Bowtie2 in local and dovetail mode which showed consistent results. BAM files were converted to BED files using the bamtobed command in the bedtools package. Coverage maps were visualised on the IGV software and reads were filtered by mapping quality (mq). Reads between 107 to 247 bp were selected for analysis to cover reads that correspond to mononucleosomes, and nucleosome dyads were peak-called using the NucleR package (v2.28)^64^ with a 1-bp resolution. Dyad maps were normalised for read number and used for further analysis.

### Fibre-seq

Fibre-seq using 5mC-methylation and nanopore sequencing, including its analysis, was performed as described previously^31^. In summary, reconstituted chromatin (1.25-1.4µg per sample) was methylated in subsequent reactions with 150 units M.*Cvi*PI and 60 units M.*Sss*I at 30°C for a total of 30 min. Methylation reactions were halted by the addition of 1% SDS, mixed with RNase A and 2 μl Proteinase K and incubated for 1 h at 50°C after which the DNA was purified by phenol:chloroform extraction. The template band was extracted from agarose gel using the Wizard® SV Gel and PCR Clean-Up System (Promega; A9282) to remove residual DNA derived from the NCPs. The template DNA concentration was assessed by Qubit fluorometer and processed into a sequencing library using the Ligation Sequncing Kit (Oxford nanopore; SQK-LSK109) and the Native Barcoding Expansion 1-12 (Oxford nanopore; EXP-NBD104) according to manufacturer instructions. The final library was loaded on a Flongle flowcell (Oxford nanopre; FLO-FLG001) and sequenced using a MinION sequencing device for 8 h. Basecalling was done by Guppy (5.0.7). After basecalling, Tombo (1.5.1) was used to resquiggle the resulting sequences to the reference genome and to subsequently calculate the methylation likelihood using the build-in 5_m_C model. The resulting files were read and processed using custom plotting commands in Python.

### Sucrose gradient sedimentation analysis of folded chromatin

Chromatin was reconstituted at a 20:1 NCP:template ratio in TEP80 buffer and the three templates were pooled. CE chromatin (-/+H5) (prepared in TEP80 as described above) was added to the template chromatin pool at a 1000:1 CE:template mass ratio to facilitate H5-redistribution right before sucrose gradient sedimentation. Sucrose gradient sedimentation was performed as described^32^. Briefly, 6-40% isokinetic sucrose gradients were prepared in TEP80, and gradients were allowed to settle overnight at 4°C. Chromatin samples were layered on the gradient and sedimentation was carried out at 41,000 rpm for 2.5 or 3 hours in a Beckman Coulter SW41 rotor. Gradients fractions were recovered by upward displacement and 40 fractions were collected from each sample. Aliquots from gradient fractions were diluted 1:5 in ddH2O and 2 μl were used for end-point PCR. Primers were designed targeting the unique junction point between different 601-601, 601-BLG or BLG-BLG fragments to enable the differentiation between fibres in the pool. Gels were imaged using the Fujifilm FLA-5100 laser scanner and densitometric analysis was undertaken using the AIDA 1D Image Analysis software. Peak-fitting was performed using PeakFit software by the Residuals procedure to determine peak width at half height.

### Large-scale chromatin preparation for SAXS

Template DNA was reconstituted at a 20:1 NCP:template ratio, without the addition of BSA, and dialysed to TEP600 buffer using 15.9 mm Visking dialysis membrane (12-14 kDa MWCO) (SLS). 50 μg of template DNA was reconstituted in a 500 μl volume per batch. Nucleosome core particles were removed by sucrose gradient sedimentation. Briefly, a 6-40% sucrose gradient was prepared in TEP600 buffer and gradients were settled overnight at 4°C. Chromatin samples were layered on the gradient and sedimentation was carried out at 41,000 rpm for 8 hours at 4°C using a Beckman Coulter SW41 rotor. Gradient fractions were recovered by upward displacement and 25 fractions were collected from each sample. Aliquots were run on a 1.2% agarose gel to determine pure template fractions. Template fractions were pooled and sucrose was removed using a PD-10 desalting column (Cytiva). Samples were then dialysed to TEP80 buffer and concentrated using Amicon® Ultra-4 centrifugal filter units after blocking with 0.1% (v/v) TritonX-100. The matched buffer was saved for background subtraction. Samples were used for SAXS at a concentration of approximately 100 ng/μl.

### Small-angle X-ray scattering (SAXS)

Small-angle X-ray scattering was performed at Diamond Light Source B21 Beamline. B21 operates in a fixed camera length configuration (3.7 m), at an energy range of 9.5-14 keV. B21 measures a resolution range from 0.0026 to 0.34 Å^-1^ using a flux of 2 × 10^12^ photons per second and is optimised for solution state SAXS experiments^65^. Purified chromatin samples were folded by the addition of purified linker H5 to template chromatin at a 1:1 H5:nucleosome ratio and incubation on ice for 30 mins. Samples were analysed by direct injection into a temperature-controlled quartz capillary cell and exposure for 3 min. Data reduction and analysis were performed using the ScÅtter3 software. Time frames were combined, excluding frames affected by radiation damage or due to a meniscus in the capillary, to give the average scattering curve for each template measurement. The average scattering from a matched buffer was then used for background subtraction. The real-space periodicities were calculated from the raw scattering data by the equation D (nm) = (2π/*q*)/10. Data was presented as reciprocal distance (nm)^-1^ as described previously^66^. A peak at q = 0.05 nm^-1^ indicates a periodicity (D) of 200 Å (20 nm).

### Transmission electron microscopy (TEM) of folded nucleosome arrays

DNA for TEM experiments was generated by digesting approximately 1 mg of plasmid DNA with *Nde*I*, Pci*I*, ApaL*I, *BspH*I, and *Eci*I overnight at 37°C shaking at 650 rpm. *Nde*I and *Pci*I digestion releases the template from the vector and generates an approximately 600 bp overhang on the end of the 5 kb fragment; while *ApaL*I*, BspH*I, and *Eci*I digestion fragments the vector backbone into fragments of less than 800 bp. The digestion was checked for completeness the following day on a 1% agarose gel and the insert was purified from the vector fragments using S-1000 chromatography as described above. The *Nde*I ends were filled with dATP (Roche) and labeled with biotin-16-dUTP (Roche) using Klenow Fragment (3’ à 5’ exo-) (NEB). The reaction was incubated at 37°C for 30 minutes and excess nucleotides were removed using the Illustra™ MicroSpin™ G-25 columns (Cytiva) and the samples were purified using phenol:chloroforom extraction and ethanol precipitation. Biotin labelling was checked using a dot blot with Streptavidin-AP (Roche).

Biotin-labelled DNA was used to reconstitute chromatin at a 20:1 NCP:template ratio in TEAP80 buffer (10 mM TEA pH 8.0, 80 mM NaCl, 0.2 mM EDTA, 0.2 mM PMSF). Chromatin quality was checked by EMSA and the concentration of the recovered template chromatin was estimated by measuring the total recovery and dividing by 20. Approximately 3 μg of template chromatin was diluted to 100 μl in TEAP80 and used for bead binding. 20 μl of the Dynabeads™ MyOne™ Streptavidin C1 beads (Invitrogen) was used for each sample. Beads were washed with TE buffer 3 times then resuspended in 100 μl TEAP80 buffer. Beads were added to chromatin in Eppendorf LoBind microcentrifuge tubes and incubated for 2 hr at 4°C with gentle rotation. For H5-redistribution, CE chromatin (+H5) (prepared in TEAP80 buffer as above) was added to the template chromatin at a 30:1 CE:template chromatin mass ratio and incubated on ice for 30 min. Chromatin was crosslinked with 0.1% (v/v) glutaraldehyde overnight at 4°C. To purify template from CE chromatin and NCPs, and to quench and remove GA, a biotin pull-down was performed on a magnetic rack and beads were washed twice and resuspended in TEP80 buffer. The template was eluted from the beads by digestion with *Xho*I (R1/R2 samples), and *Eco*RV-HF (601 sample) in 0.1× Cut Smart Buffer (NEB) at 25°C overnight with gentle rotation, and the beads were removed after incubation on a magnetic rack. The supernatant containing the template chromatin was dialysed to TEP80 buffer for 4 hr at 4°C using the Tube-O-DIALYZER™ mini dialysis system (50 kDa MWCO; G-BioSciences). Chromatin was coated in benzalkonium chloride (BAC) by incubating in 2×10^−4^ % (v/v) BAC for 1 hour at room temperature, and 5 μl of sample (∼0.5 - 1 ng/μl) was adsorbed onto carbon/formvar 200 mesh grids (TAAB technologies) for 5 minutes then washed with ddH2O two times. The grids were blotted gently on filter paper then placed on a 50 μl drop of 90% (v/v) ethanol for 3 seconds and blotted again. Grids were transferred to a filter paper and left to dry. To enhance contrast a 3 nm layer of platinum was evaporated onto the grids using the Leica EM ACE600 High Vacuum Sputter Coater at a 7° angle. Grids were imaged on the JEOL JEM-1400Plus Transmission Electron Microscope at 15,000-25,000× magnification.

TEM Analysis was performed using the MountainSPIP analysis software (v8, Image Metrology) using a custom automated pipeline. Briefly, TEM images of individual fibres were selected and processed by image smoothing using a 25-point Gaussian filter. The resulting images were segmented to threshold fibres (threshold set at 5 grey scale levels away from the mean plane) with area larger than 2000 nm^2^. The mean diameter, length, and skeleton lengths of each fibre were quantified. Cluster quantification was performed in FIJI (ImageJ). A region of interest was drawn over individual fibers, and an automated macro was applied to process the images. The macro included a Gaussian blur to reduce noise, followed by Phansalkar local thresholding and watershed segmentation. Particle analysis was then used to identify and count clusters within each segmented fiber.

### Atomic Force Microscopy (AFM) imaging and data analysis

Samples containing Widom 601 25-mers, in the absence of linker histone, and with or without glutaraldehyde crosslinking were deposited from aqueous buffer (10 mM Tris-HCl, pH=7.5, 0.2 mM EDTA, 0.2 mM PMSF, 80 mM NaCl) on poly-L-lysine (M_W_ = 1000-5000) coated mica following a protocol described previously^67^. Briefly, poly-L-lysine coating was undertaken by drop casting a 0.01% (w/v in autoclaved milliQ water) solution onto freshly cleaved mica for 30 seconds, followed by gentle rinsing with 20 mL milliQ water, and drying using N_2_ gas. The coated mica discs were immediately used for sample deposition (15 seconds) followed by rinsing with milliQ water (20 mL) and drying under a stream of N_2_ gas.

Samples were imaged using a commercial JPK Nanowizard 4 XP AFM (Bruker) in tapping mode in air, immediately after preparation. SSS-NCHR cantilevers (Nanosensors) were used and excited near their resonance frequency (∼300 kHz) and at ∼85% of the free air amplitude. Imaging speed and feedback parameters were optimized to allow stable imaging.

AFM topographs were processed and analyzed using SPIP software (v6.1., Image Metrology). Processing involved background subtraction and image erosion using blind tip reconstruction. The particle analysis module in the SPIP was used to quantify nucleosome cluster volumes (height threshold 0.5 nm).

### Single-molecule force spectroscopy (SMFS) with magnetic tweezers

DNA for magnetic tweezers experiments was prepared by digestion as described above with *Nde*I and *Pci*I and vector fragmentation. The *Nde*I ends were filled with dATP (Roche) and labeled with biotin-16-dUTP (Roche) using Klenow Fragment (3’ à 5’ exo-) (NEB). The reaction was incubated at 37°C for 30 minutes and excess nucleotides were removed using the Illustra™ MicroSpin™ G-25 columns (Cytiva). The *Pci*I end was filled in with dATP, dCTP, dGTP (Roche) and labelled with DIG-11-dUTP (Roche) using Klenow Fragment (3’ à 5’ exo -) (NEB). The reaction was incubated for 2 hours at 37°C and the samples were purified using phenol:chloroforom extraction and ethanol precipitation. Samples were stored at −20°C, and labelling was checked using a dot blot with Streptavidin-AP (Roche) or Anti-Digoxigenn-AP, Fab fragments (Roche).

Home-made flow cells were prepared by cutting flow channels of approximately 10 × 40 × 0.4 mm on parafilm sheets and attaching them to a glass coverslip followed by mounting on a plastic frame with holes for sample injection. Prior to assembly the glass coverslip was cleaned by sonication for 15 mins in isopropanol and treated with 0.1% (v/v) nitrocellulose in amylacetate to retain a hydrophobic surface on the glass slide. Flow channels were injected with 50 μl of 10 μg/ml Anti-DIG (Roche) (diluted in PBS) and incubated for 2 hr at 4°C. Channels were blocked by the addition of 50 μl of 4% (w/v) BSA (diluted in PBS) and incubated overnight at 4°C. Flow cells were stored at 4°C for up to one week.

Biotin+DIG-labelled template DNA was reconstituted at a 20:1 NCP:template ratio in TEP80 buffer. Chromatin quality was checked by EMSA and the concentration of the recovered template chromatin was estimated by measuring the total concentration and dividing by NCP ratio used. Flow channels were washed with measurement ESB+ buffer (10 mM HEPES pH 7.6, 100 mM KCl, 10 mM NaN3, 0.1% (v/v) Tween-20, 0.2% (w/v) BSA, 2 mM MgCl_2_) prior to the injection of chromatin. Typically 5 ng of template chromatin was diluted to 50 μl in ESB+ buffer and injected in the flow cell channel then incubated for 15 mins at room temperature to allow DIG-attachment to the glass surface. Channels were washed with ESB+ buffer to remove donor NCPs, and 0.05 μl of Dynabeads™ M-270 Streptavidin (Invitrogen) (diluted in 50 μl ESB+ buffer) was injected and incubated at room temp for 15 min to facilitate binding of the biotinylated DNA end to beads. 250 ng of CE chromatin (-/+H5) (diluted to 50 μl in ESB+ buffer) was added for a 50:1 CE:Template chromatin mass ratio and incubated at room temperature for 15 min to promote H5-redistribution. Flow-cells were washed with 50 μl ESB+ buffer to wash CE chromatin, and then incubated for 15 min at room temperature before the initiation of magnetic pulling to equilibrate the temperature of the flow channel. All flow-cell injection steps were performed gently to minimise bead detachment due to drag forces.

A custom-built multiplexed magnetic tweezers system was used to manipulate chromatin fibres and measure the extension of fibres under increasing force^68^. A force ramp trajectory was applied increasing from 0 pN to 43 pN, followed by reversal of the magnet trajectory to decrease force. For fibres reconstituted with truncated tails, a force ramp trajectory was applied increasing from 0 pN to 6.25 pN, followed by reversal of the magnet trajectory to decrease force. A second force trajectory from 0 to 43 pN was then applied to the same field, followed by force reversal. The force exerted was calculated using a double exponential function calibrated prior to the experiment. The change in height of the magnetic beads (and therefore the extension of chromatin molecules) was measured in real time at a frame rate of 30 Hz with a digital camera (CMOS Vision Condor).

Analysis was performed using custom software written in LabView^50^. Chromatin traces that did not show significant extension, reflecting stuck beads, were discarded. To correct for off-set in the z-direction, the force-extension profile for individual fibres was aligned at the most extended part of the profile (F > 35 pN), where the nucleosomes in the fibre are expected to fully unwrap to a worm-like chain (WLC) of 5590 bp with a persistence length of 50 nm and a stretch modulus of 1000 pN. A heatmap was then generated for all chromatin force-extension traces within the same condition.

### NucHoP model for chromatin

We have developed NucHoP (Nucleosome-resolved Higher order Polymer) model for chromatin. A course-grained molecular dynamics simulation modelling approach was used, whereby groups of atoms and molecules were represented by spherical ‘beads’ or collections of beads (Extended data 10). The geometry of the nucleosome was obtained from the canonical 601 nucleosome crystal structure (PDB:8JBX)^69^; a set of beads is positioned to provide the steric interactions of the cylindrical nucleosome core particle (core histones and wrapped DNA). Nucleosome core particles are treated as rigid bodies (i.e., there are no internal motions of the histone proteins, and the DNA cannot unwrap from the core—unwrapping is likely less relevant for chromatin fibres with linker histone^45^), and are connected by chains of beads representing DNA. The length of the DNA controls the relative equilibrium orientation of neighbouring nucleosomes. We use the ‘twistable elastic polymer’ DNA model which we previously developed^70^; in this model the position and orientation of each bead is tracked, with phenomenological potentials providing an energy cost for bending and twisting, and a bond potential enforcing chain connectivity (Extended data 10, left of centre panel). Each DNA bead represents 7 bp, but some bonds between beads can be shortened to allow setting of linker lengths at base-pair resolution. The bending and torsional stiffness of the chain is matched to known average values for DNA^71,72^..

An attractive inter-nucleosome interaction potential is set to provide different attraction strengths for nucleosomes in different relative orientations (face-to-face interactions are strongest, then face-to-side, etc.). These are matched to published potentials obtained from a more detailed nucleosome simulation model^73^ (parametrised from atomistic details and itself found to be consistent with nucleosome DNA origami experiments^74^). This is achieved in our model by specifying an interaction potential between each of the beads within a pair of nucleosome cores such that the sum of these interactions gives the desired internucleosomal interaction which depends on the separation and relative orientation of the cores (Extended Data 10, rightmost panel). Our model matches the shape and relative strength of interactions in different orientations reported in previous studies^73^; the overall strength of nucleosome-nucleosome stacking attraction (the depth of the face-to-face potential well in the model) depends strongly on salt concentration and has not been well characterised, with different values obtained via different experimental methods^22,74,75^. We treat this as a parameter in the model, choosing a value which is just strong enough to give compaction of the 601 fibres (mapping the interaction energy scale from simulation to real units is not straightforward, but in a simulation of two mono-nucleosomes, this leads to a nucleosome stacking residency time which is approximately consistent with the 2.69 k_B_T energy previously described^73^ as being relevant for physiological salt conditions).

Linker histone is not modelled explicitly, but its effect on nucleosome structure is included in two ways: as revealed in cryoEM experiments^45^, presence of linker histone alters the equilibrium opening angle of the exiting DNA and effectively stiffens that DNA to reduce fluctuations in opening angle. Fibres can be generated with any pattern of DNA linker lengths (which are then fixed for the duration of a simulation), and fibre configuration dynamics evolved using an implicit solvent molecular dynamics scheme (Langevin dynamics) using the LAMMPS software^76^. Each simulation was first run for a time equivalent to 7.2 ms to allow the fibre to fold into a configuration representative of equilibrium; then a ‘production simulation’ of duration equivalent to 36 ms was run, with configurations saved at intervals of 0.36 ms. For the simulations presented in Fig. 4a, configurations were saved more frequently to allow tracking of measured quantities through time.

For 601, nucleosomes were separated by 50 bp. For R1 and R2 fibres, the ‘NucPosSimulator’Click or tap here to enter text. software^77^ was used to obtain sets of nucleosome positions consistent with the MNase-seq data for each template. To obtain the ‘phased’ nucleosome positions, NucPosSimulator was used in a mode where is finds a single set of nucleosome positions which best fit the MNase-seq; these are ‘phased nucleosome arrays’, where the nucleosome positions are the same for each simulated fibre. To obtain the ‘unphased’ nucleosome positions, NucPosSimulator was instead used in a mode where it generated multiple different sets of positions consistent with the relevant MNase-seq data. These are ‘unphased nucleosome arrays’, which capture fibre-to-fibre nucleosome position variation. All simulated fibres contained 25 nucleosomes.

As detailed in the main text, force-extension simulations are performed by attaching one end of a fibre to an array of beads to which a force is applied, while the other end of the fibre is held fixed (Fig. 5e). The array of beads can rotate in the *x*-*y* plane, such that the fibre is not under any torsional constraint, but its motion is constrained to be in the *z*-axis direction (the direction in which the force is applied). The magnitude of the force required to, e.g., “unstack” a pair of nucleosomes will depend on the strength of the stacking interactions which, as discussed above, is a parameter in our model. This, together with the complications in mapping simulation energy scales to real energies, means that the values of the force are not directly comparable between simulations and SMFS experiments.

### Chromatin polymer model simulation measurements

To quantify the size of a fibre the radius of gyration (using the standard definition) is calculated for each given configuration using the positions of all nucleosome core beads. The shape anisotropy is a function of the eigenvalues of the gyration tensor, and this would have a value of 0 if the nucleosome bead positions were distributed spherically symmetrically and 1 if the positions were in a line. To calculate the stacking fraction, we define two nucleosomes faces as being stacked if the beads which sit in the centre of each face have a separation within a threshold distance of 8.4 nm, and the orientation of each nucleosome deviates by no more than 45 degrees from the line connecting their centres.

To quantify the accessibility, for each given DNA bead, the size of the largest sphere which could fit around that bead without interacting with any part of a nucleosome core was calculated (Extended Data 13). To obtain the accessibility score for a given fibre configuration the size of this sphere for all DNA beads within the linkers was averaged.

To obtain interaction maps (simulated ‘Micro-C’), from the positions of the DNA and nucleosome beads, we used an interpolation scheme to assign a position to each base-pair of the DNA. We calculated the separation between each pair of base-pairs, and counted it as a ‘contact’ if this was less than 3 nm. We constructed the contact map at 10 bp resolution by considering the fraction of configurations within a simulation where the two base-pairs are in contact. These maps are therefore at a much higher resolution that would be possible to obtain experimentally.

The nucleosome positioning irregularity (which does not vary during a simulation) is defined as the mean squared deviation of the linker lengths away from 50 bp; i.e., it is given by

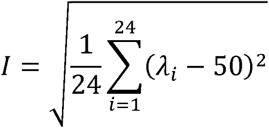

where the sum runs over all over the linker DNA segments where *λ_i_* is the length of the linker between nucleosomes *i* and *i*+1. The irregularity therefore has units of bp.

