## Supplementary Figures for "Irregular nucleosome positioning governs a crystalline to liquid-like phase transition and tunes chromatin accessibility"

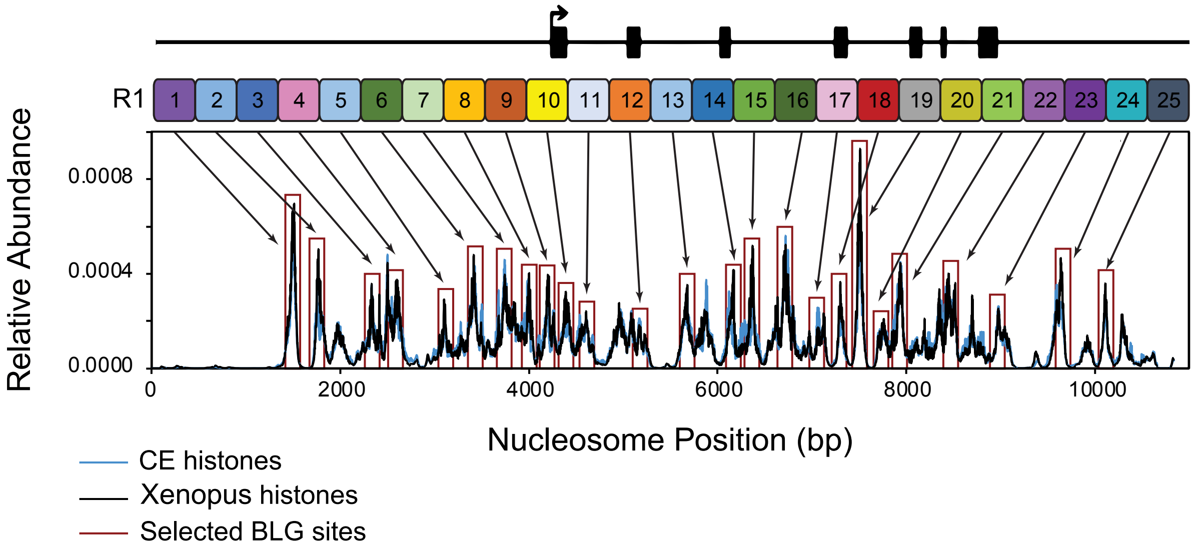


AlBawardi et al., Extended Data1

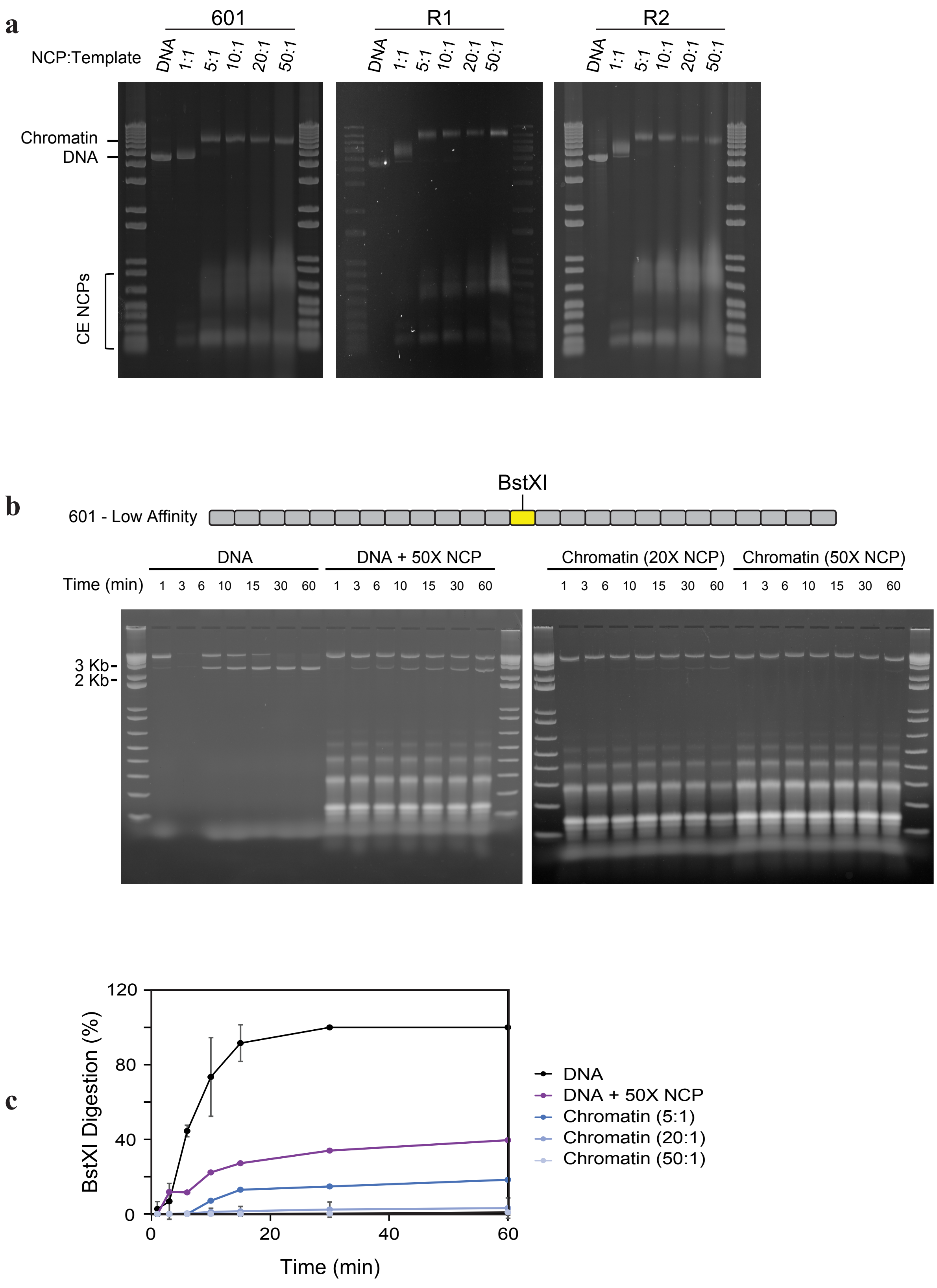


AlBawardi et al., Extended Data 2

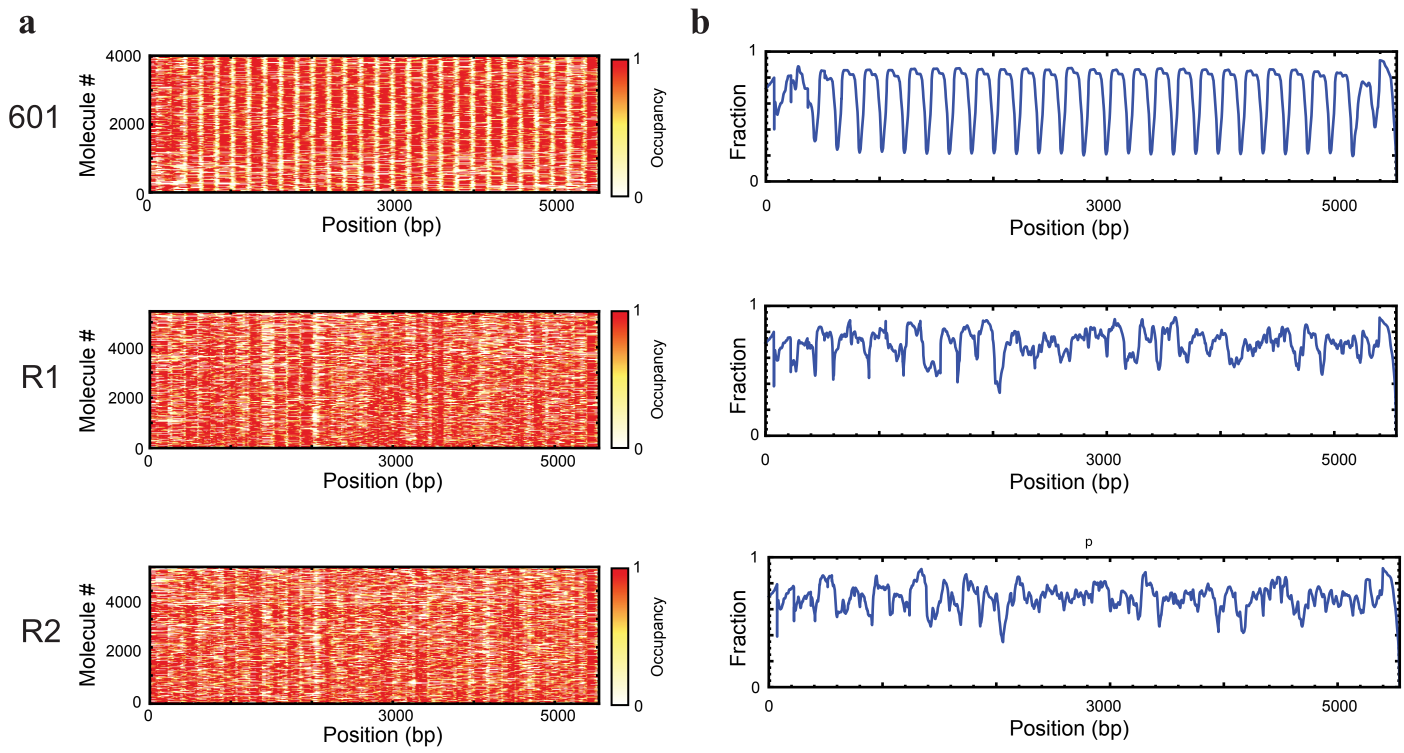


AlBawardi et al., Extended Data 3

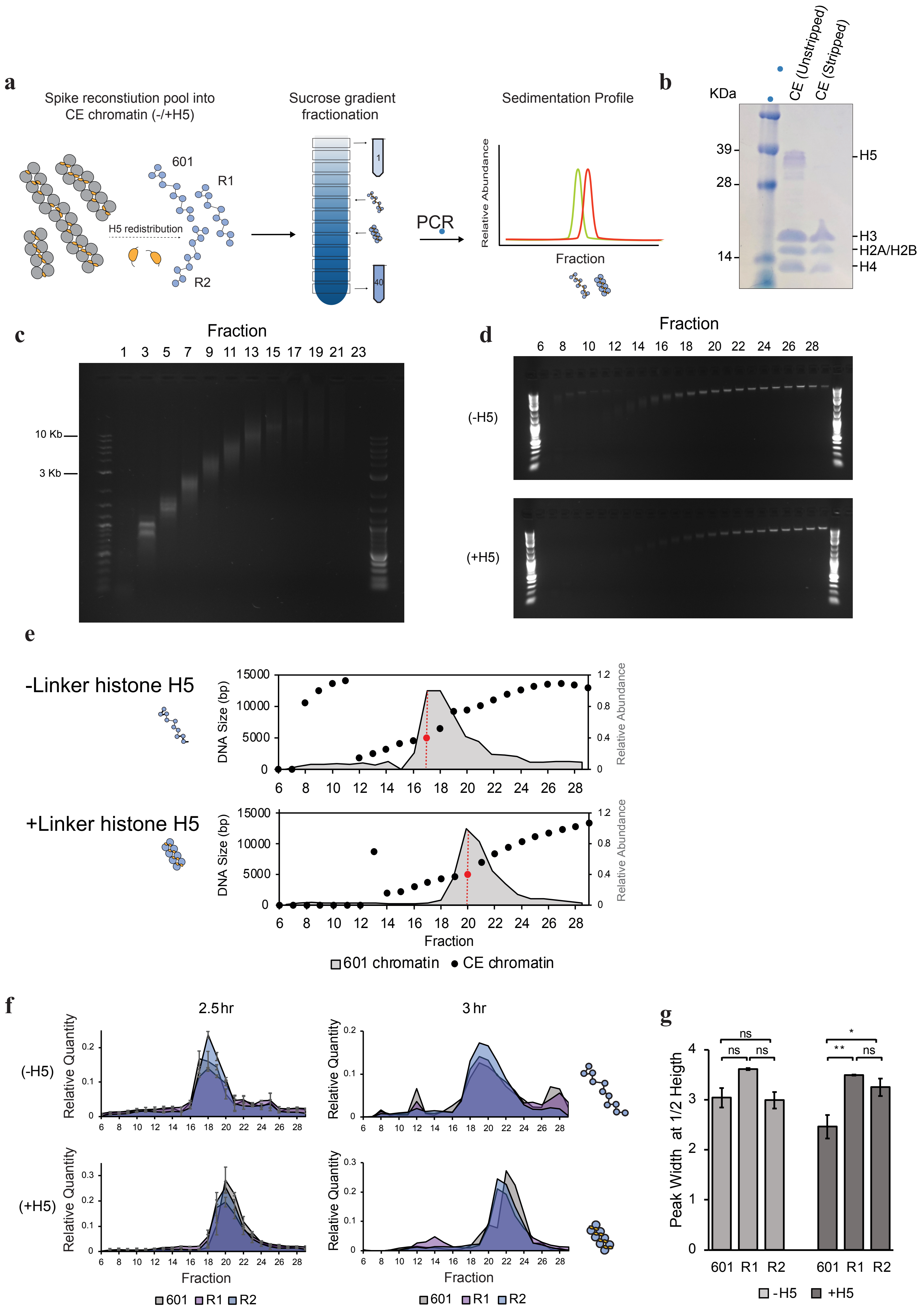


AlBawardi et al., Extended Data 4

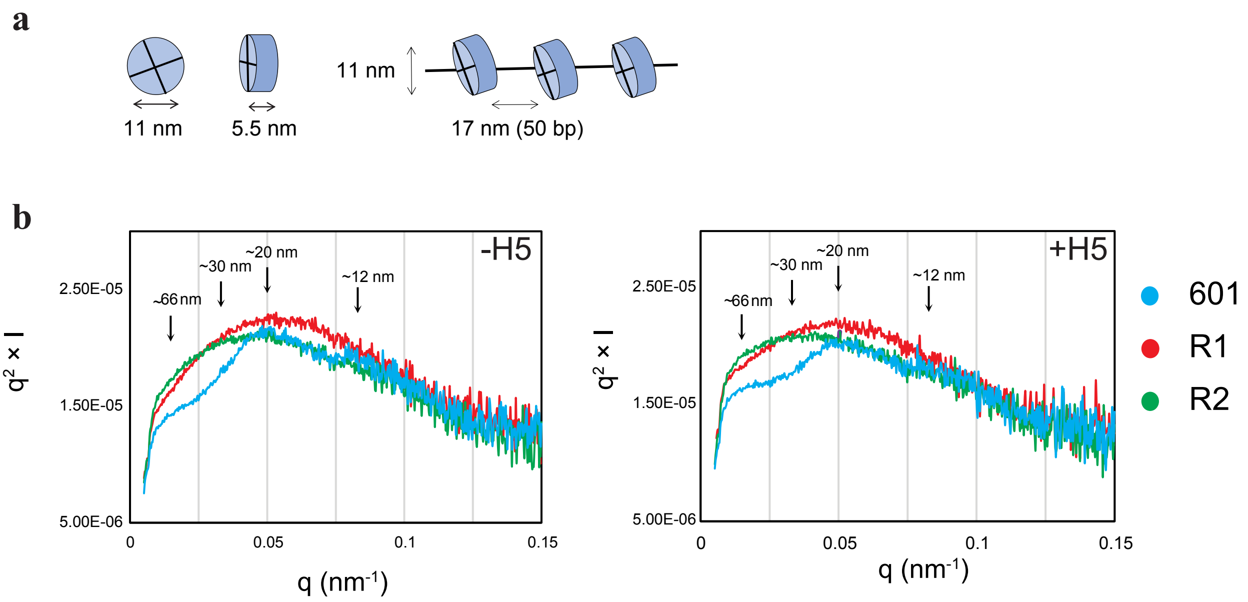


AlBawardi et al., Extended Data 5

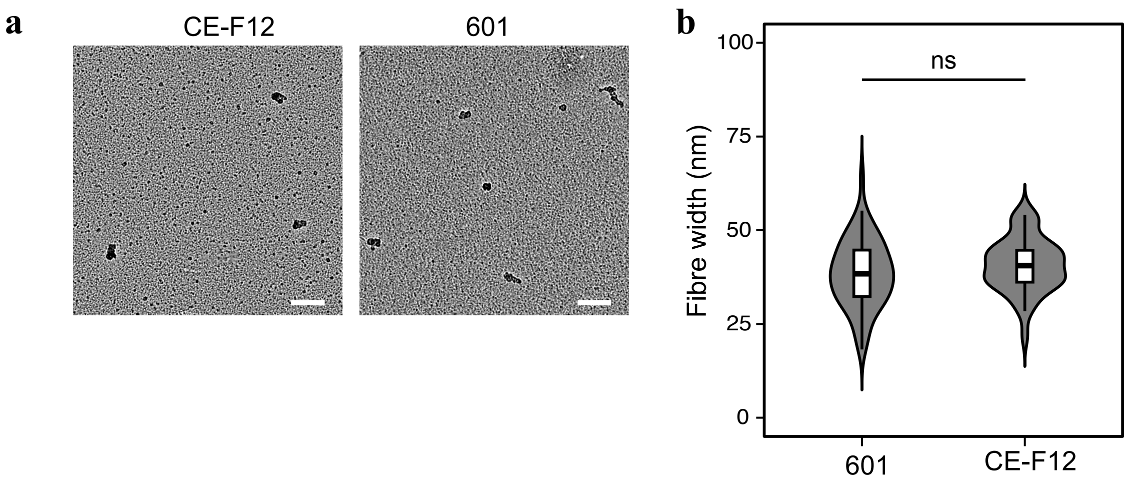


AlBawardi et al., Extended Data 6

601

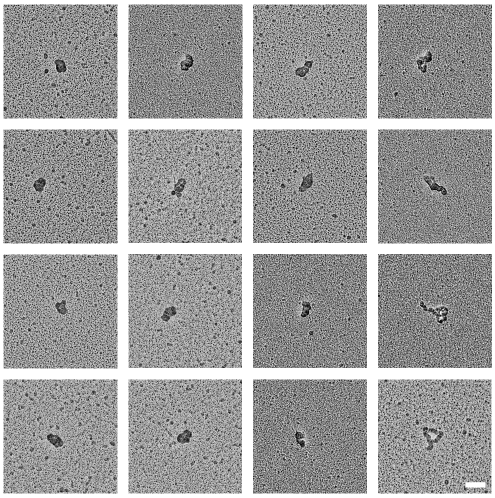

R1

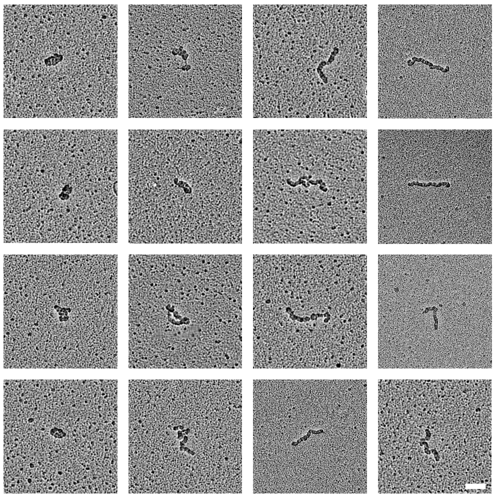

R2

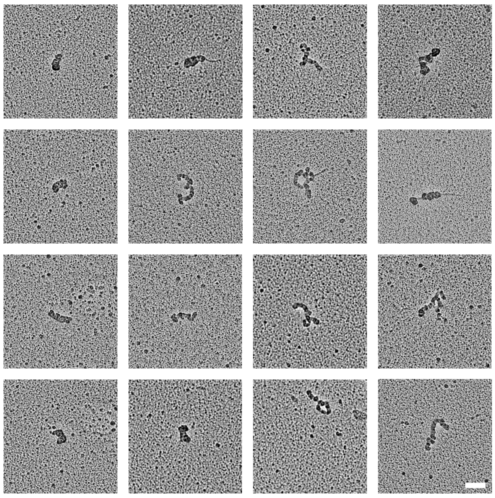

AlBawardi et al., Extended Data 7

**a**

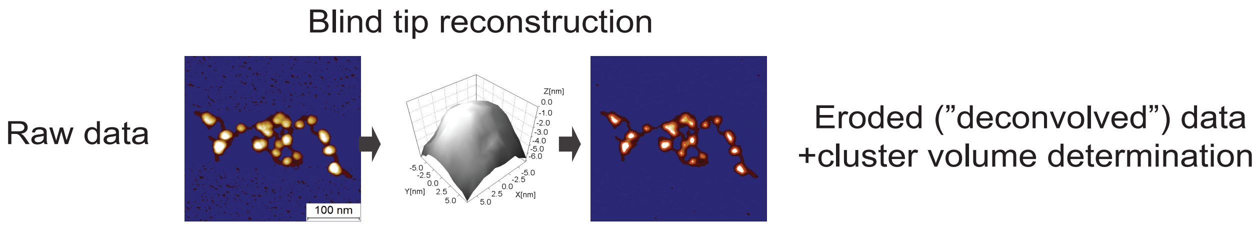


**b**

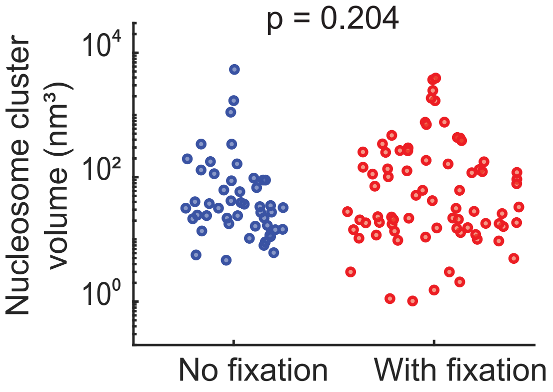


AlBawardi et al., Extended Data 8

**a**

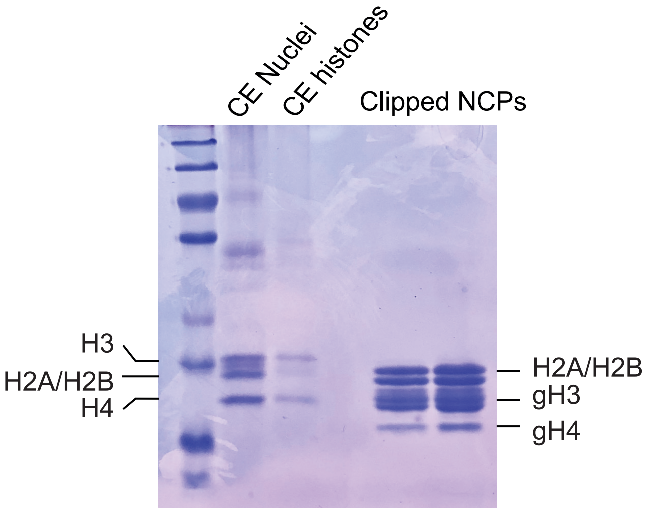


**b**

| | Expected mass (Da) | Detected mass (Da) |
|---|---|---|
| H2A | 13940 | 13650 |
| H2B | 13922 | 13852 |
| H3 | 15256 | 12740 |
| H4 | 11367 | 8474 |

**c**

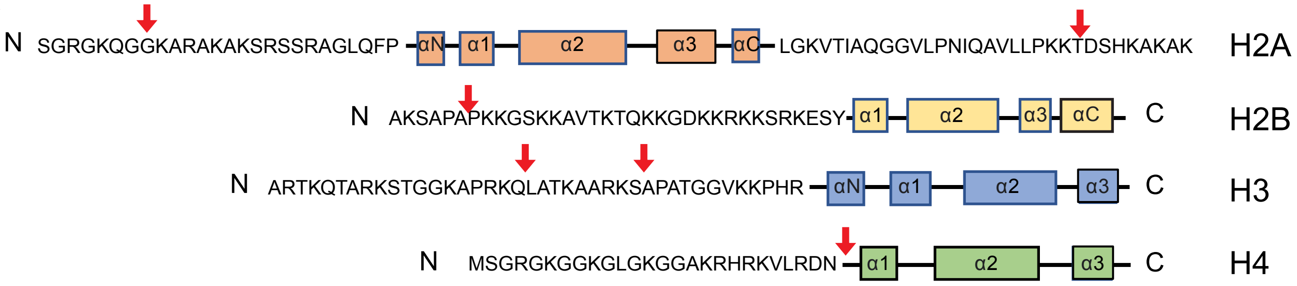


AlBawardi et al., Extended Data 9

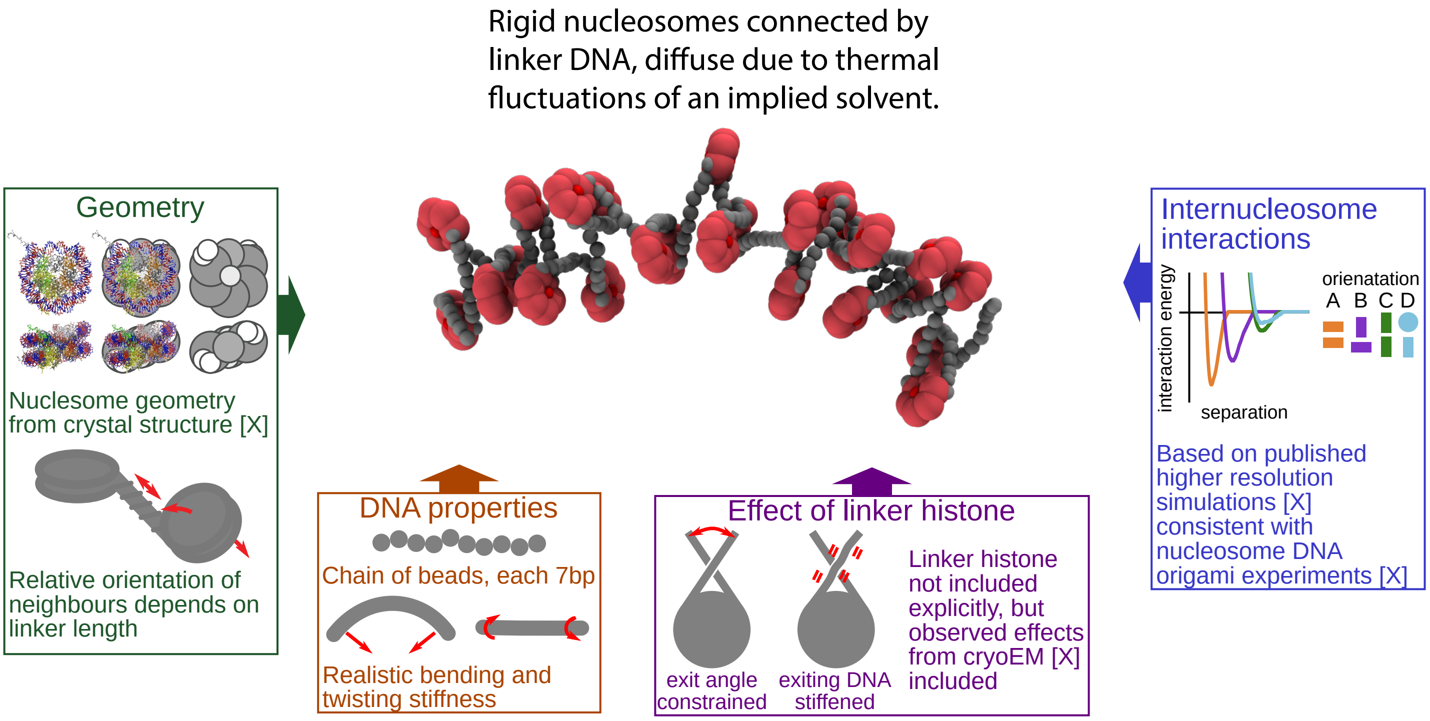


AlBawardi et al., Extended Data 10

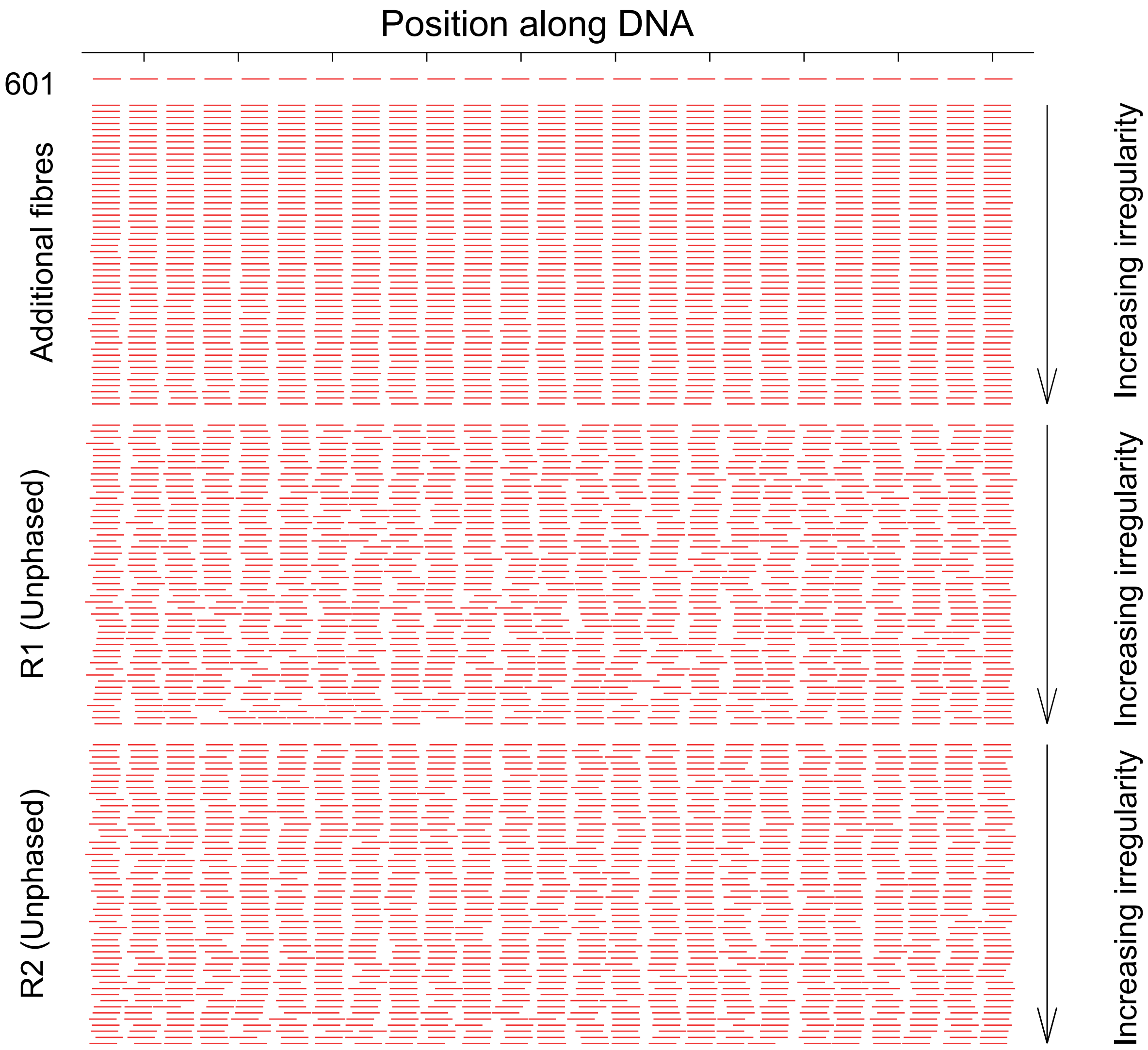


AlBawardi et al., Extended Data 11

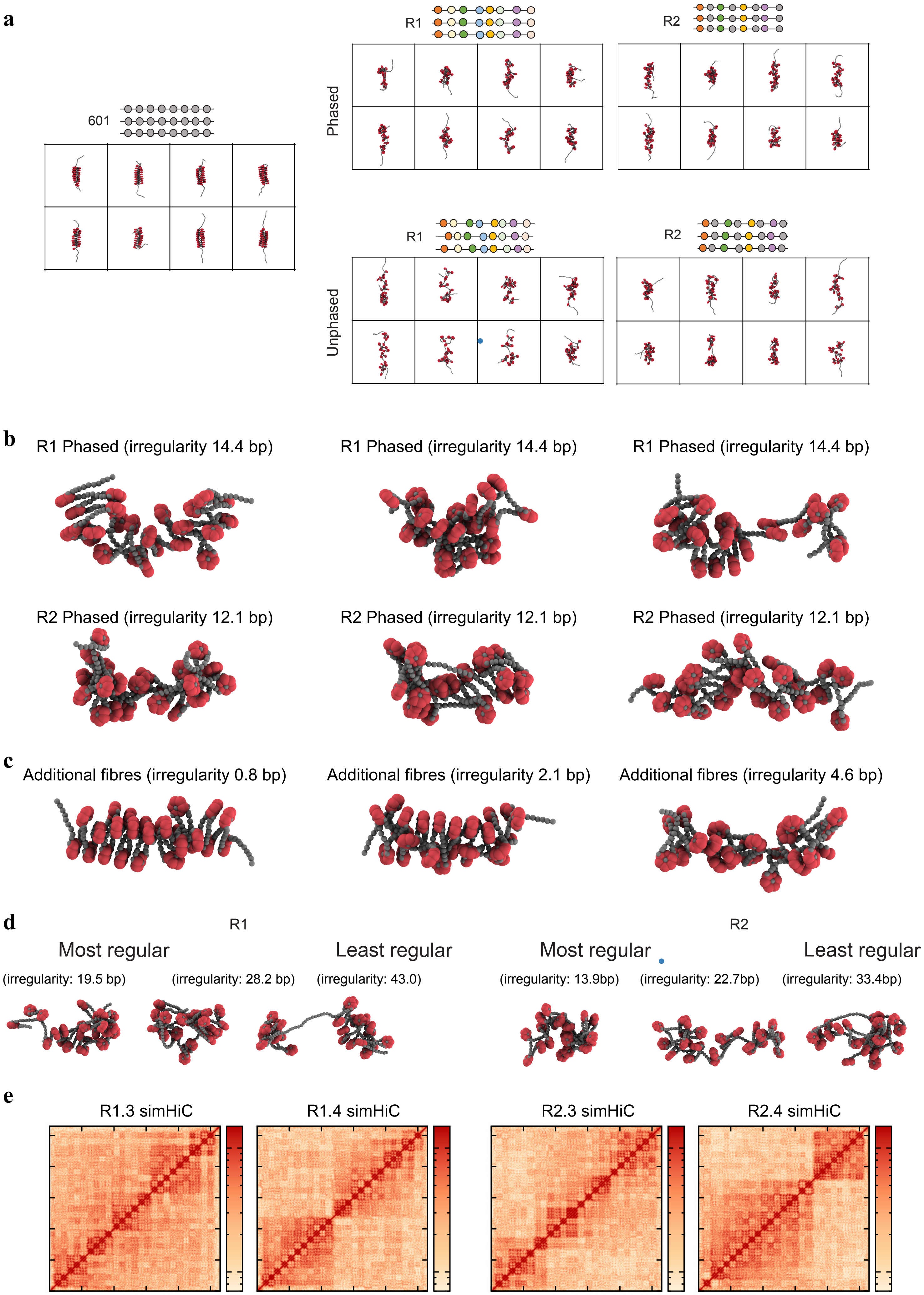


AlBawardi et al., Extended Data 12

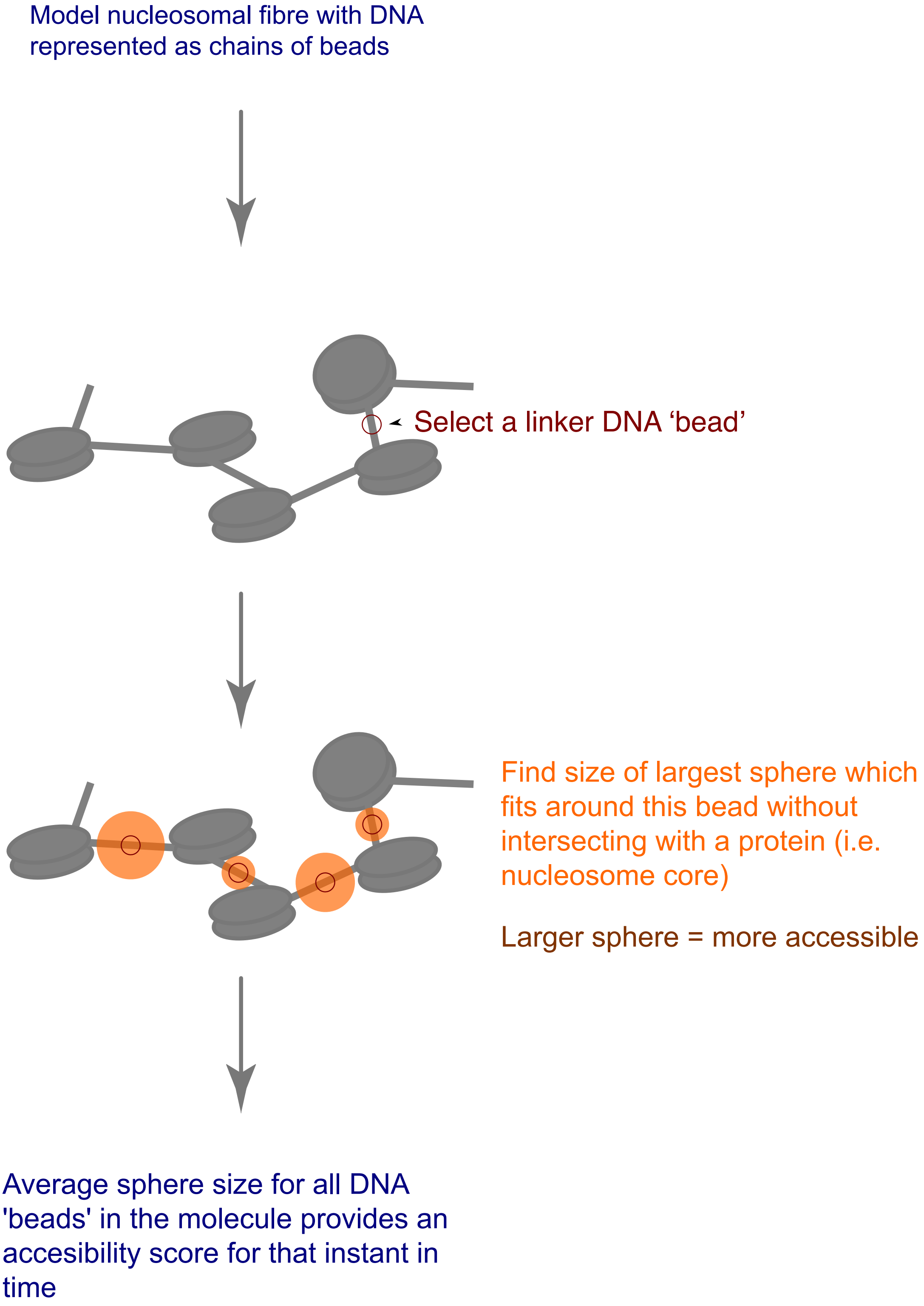


AlBawardi et al., Extended Data 13

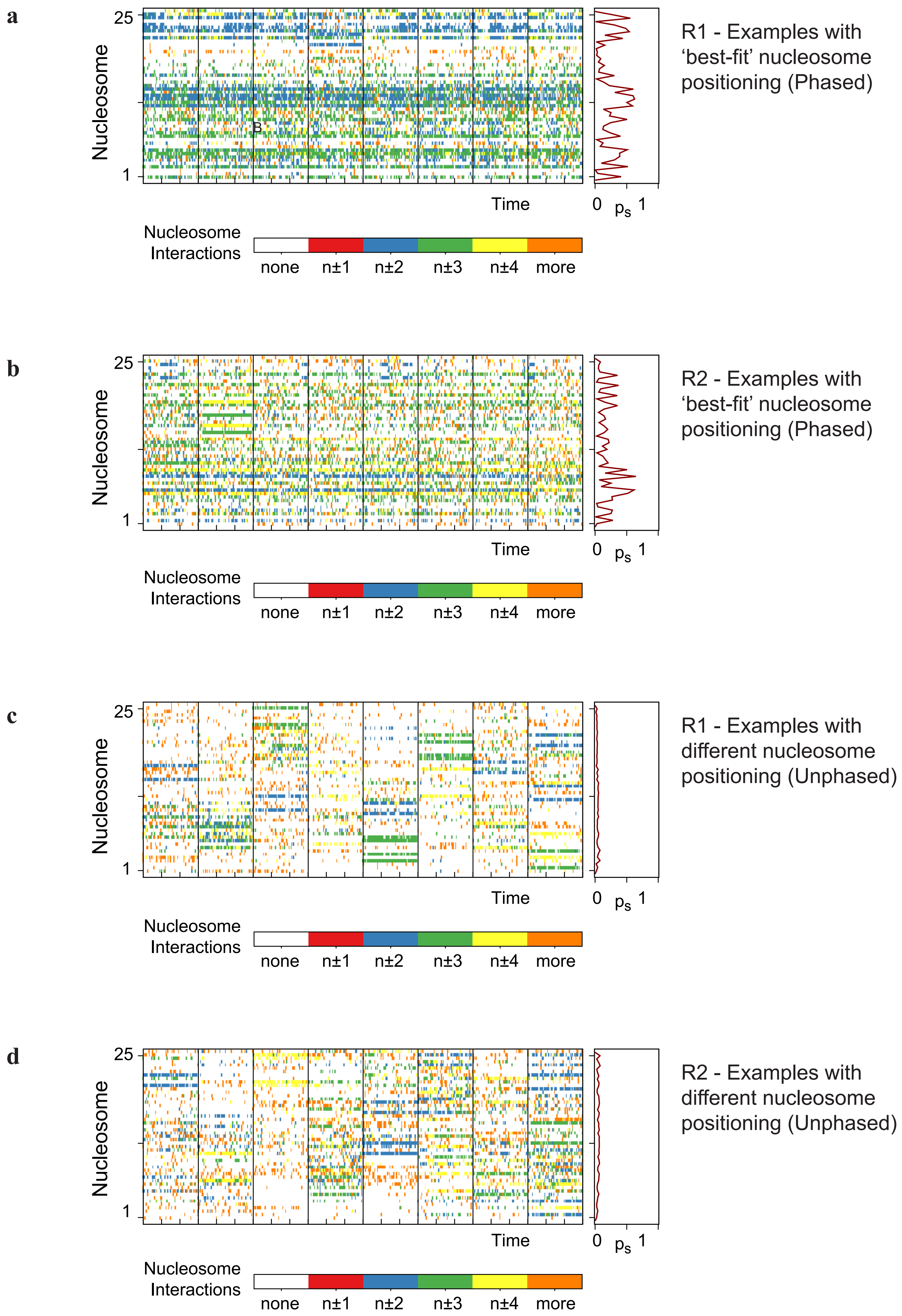
a
b
c
d
25
1
Nucleosome
Time
0
$p_s$
1
Nucleosome Interactions
none
n±1
n±2
n±3
n±4
more
R1 - Examples with 'best-fit' nucleosome positioning (Phased)
R2 - Examples with 'best-fit' nucleosome positioning (Phased)
R1 - Examples with different nucleosome positioning (Unphased)
R2 - Examples with different nucleosome positioning (Unphased)

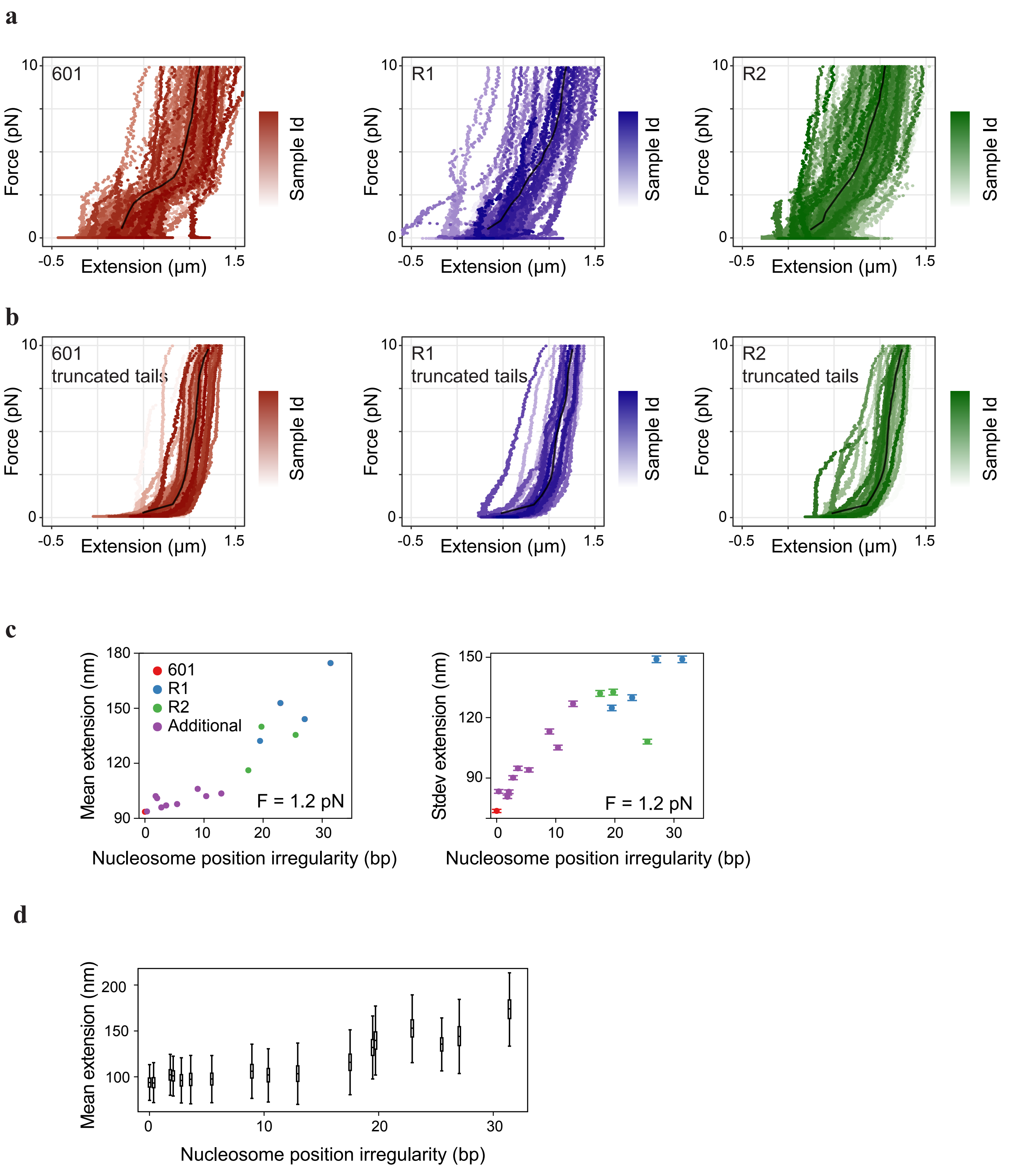


AlBawardi et al., Extended Data 15
